# Toward a unified theory of microbially-mediated invasion

**DOI:** 10.1101/2024.01.18.576278

**Authors:** Maria M. Martignoni, Jimmy Garnier, Rebecca C. Tyson, Keith D. Harris, Oren Kolodny

## Abstract

Biological invasions pose major ecological and economic threats, and extensive research has been dedicated to understanding and predicting their dynamics. Most studies focus on the biological invasion of single species, and only in recent years has it been realized that multi-species interactions that involve native and invasive host species and their microbial symbionts can play important roles in determining invasion outputs. A theoretical framework that treats these interactions and their impact is lacking. Here we offer such a framework and use it to explore possible dynamics that may emerge from the horizontal sharing of native and non-native microbial symbionts among native and non-native host individuals and species. Thus, for example, invasive plants might benefit from native mycorrhizal networks in the soil, or might be particularly successful if they carry with them parasites to which competing native hosts are susceptible. On the other hand, invasion might be hindered by native parasites that spread from native to invasive individuals. The mathematical framework that we lay out in this study provides a new mechanistic, cohesive, and intuition-enhancing tool for theoretically exploring the ways by which the subtleties of the relationships between hosts and microbes may affect invasion dynamics. We identify multiple pathways through which microbes can facilitate (or prevent) host invasion, microbial invasion, and the invasion of both hosts and their co-introduced microbes. We disentangle invasion outcomes and highlight modalities of ecological dynamics that have so far not been considered in invasion biology. Our work sets the foundations for invasion theory that includes a community-level view of invasive and native hosts as well as their microbial symbionts.

## 1 Introduction

Biological invasion is often regarded as invasion by a single species. However, organisms live in symbiosis with a rich and diverse collection of microbial symbionts that are an essential component of their host’s fitness and reproductive success (Fitzpatrick et al., 2020; Compant et al., 2019). Microbial communities can encompass mutualistic, commensal, and parasitic (or pathogenic) symbionts that benefit or harm the host to different extents. The fitness of these microbial symbionts, in turn, depends on their association with host partners, such that the fates of hosts and symbionts are intrinsically linked, and so are the possibilities for hosts and symbionts to successfully establish and persist in a new environment. For example, the formation of novel associations between invasive plants and pre-existing native mycorrhizal networks can facilitate the establishment of an introduced host population and its expansion into a new range (Dawkins and Esiobu, 2016; Parepa et al., 2013; Moeller et al., 2015). The spread of introduced symbionts can also be facilitated by native hosts (Díez, 2005; Wolfe and Pringle, 2012; Dickie et al., 2016), where the spread of introduced pathogens may cause the emergence of disease in native species and lead to their competitive exclusion (Panzavolta et al., 2021; Santini et al., 2013; Grünwald et al., 2012; Schuchert et al., 2014). In other cases, host-symbiont interactions can provide resilience to native communities. For instance, host invasiveness may be decreased by a reduction in the abundance of their mutualistic symbionts (Zenni and Nuñez, 2013; Catford et al., 2009), or by the transmission of pathogenic agents from native to invasive hosts (Mordecai, 2013; Flory and Clay, 2013).

Awareness of the importance of host-symbiont associations in dynamics of host and microbial invasion is growing rapidly, and so are attempts at developing conceptual frameworks to understand the impact of parasitic or mutualistic host-symbiont associations on invasion outcomes (Dickie et al., 2017; Amsellem et al., 2017; Dunn and Hatcher, 2015; Médoc et al., 2017; Mitchell et al., 2006; Nuñez et al., 2009; Dickie et al., 2017). However, more studies are needed to fully understand the impact that multi-species interactions that involve native and invasive species and their microbial symbionts may have on invasion dynamics. From a theoretical point of view, ecological population modelling has mostly studied the range expansion and invasion dynamics of either parasite (White et al., 2018; Gubbins et al., 2000), or host populations (Lewis et al., 2016), rarely accounting for the role that host-symbiont feedback can have on the invasion dynamics. Even in the few instances in which these feed-backs were considered (Bever et al., 1997; Yamauchi et al., 2011; Jack et al., 2017; Martignoni and Kolodny, 2023; Kandlikar et al., 2019), studies mostly unilaterally discussed how diversity and fitness of the host population can be mediated by microbial communities and not vice-versa. Host and symbiont fitness is interdependent: symbionts can influence the growth and coexistence of host populations, and changes in host populations will, in turn, influence fitness and diversity in symbiont communities, all of which can affect competitive dynamics between hosts and between symbionts.

Theoretical insights into the stability and resilience of mutualistic and parasitic communities against invasion have been provided by network theory. These approaches have mainly focused on relating community invasiveness to network properties, such as nestedness, or connectivity (Campbell et al., 2012; Bastolla et al., 2009; Rohr et al., 2014; Vacher et al., 2010), where network structure is based on trait-based approaches considering how trait differences and similarities in the invaded and invading communities may affect interaction strength (Minoarivelo and Hui, 2016a; Hui et al., 2016; Minoarivelo and Hui, 2016b; Schleuning et al., 2015; Runghen et al., 2021). These studies can help identify common features of mutualistic or host-parasite networks, and test hypotheses regarding the relationship between structure and community dynamics (Valdovinos, 2019; Bascompte and Olesen, 2015). However, particularly for the case of mutualistic networks, analyses have been performed on communities of free-living species, such as pollination or seed-dispersal mutualisms (Bascompte and Jordano, 2013), whose dynamics may differ from what is observed when microbial symbionts are obligate mutualists. Additionally, theoretical results are largely based on Lotka-Volterra equations that may lead to inaccurate predictions due to their unrealistic biological assumptions, such as linear positive effects of mutualistic interactions and unlimited growth (Holland, 2015; Gibbs et al., 2023). Thus, the development of mathematical models that integrate biologically relevant mechanisms, such as density-dependent (instead of linear) positive effects of mutualism, while maintaining the necessary simplicity to allow analytical tractability, is key to providing a predictive understanding of the dynamics of ecological communities of hosts and their symbionts (herein referred to as ‘host-symbiont communities’), and their potential to invade or be invaded.

We develop a mathematical framework to explore the possible invasion dynamics occurring when novel host-symbiont associations can form between native and invasive species. Our model accounts for key features of host-symbiont interactions including, critically, host-symbiont interdependent fitness and density-dependent resource exchange mechanisms, where a continuum of host-symbiont interactions ranging from mutualistic to parasitic is considered. Despite their opposite effects on host fitness, invasion dynamics driven by mutualistic or parasitic host-symbiont interactions may present similar interaction motifs (Dickie et al., 2017). For instance, the formation of mutualistic associations between invasive hosts and native mycorrhizal fungi (i.e., an association that would benefit invasive hosts) and the transmission of parasites to native hosts (i.e., an association that may weaken native hosts) may lead to similar outputs, namely providing a competitive advantage to invasive hosts and increasing their invasion success. Our model allows for the theoretical characterization of these multiple invasion pathways within a single framework, wherein native host-symbiont community is confronted by an invasive host-symbiont community. Our framework has unifying features that allows us to catalogue the impact and diversity of host-symbiont interactions observed experimentally and organise them in a methodological manner that can help us establish connections between similar invasion motifs and, at the same time, enhances our mechanistic understanding of biological invasion mediated by host-symbiont associations.

We will describe multiple ways by which host-symbiont interactions can facilitate or prevent host invasion, symbiont invasion, or co-invasion, defined here as the simultaneous invasion of an introduced host population and its symbionts. We portray the model in terms that correspond best to symbioses for which microbes are primarily external to their hosts (e.g., plant-microbial symbioses), because these have been studied and described (Dickie et al., 2017; Bever et al., 2010). However, the same principles may hold for systems in which the microbes are internal or partially internal to their hosts (e.g., gut or coral microbiomes, Pettay et al. (2015); Chiarello et al. (2022); Goedknegt et al. (2017)).

## 2 Model and Methods

### 2.1 Mathematical framework

To investigate the dynamics of host-symbiont communities we develop a consumer-resource model for mutualism (Holland and DeAngelis, 2010), similar to that presented by Martignoni et al. (2020). We consider a native host population with biomass *p*_*n*_ and its associated native symbiont community with biomass *m*_*n*_. Hosts and symbionts interact by exchanging resources necessary for each other’s growth. For example, in the case of the mycorrhizal symbiosis, the host plant provides synthesized carbon in the form of sugars (e.g., glucose and sucrose) to its associated mycorrhizal fungi, and receive necessary nutrients such as phosphorus, nitrogen, or water in return (Smith and Read, 2010). The transfer of resources from hosts to symbionts increases symbiont biomass and decreases host biomass, as described by the *F*_*m*_ functions below. The transfer of resources from symbionts to hosts increases host biomass and decreases symbiont biomass, as described by the *F*_*p*_ functions below. We consider hosts to be facultative mutualists, and capable of some growth in the absence of the symbionts (with intrinsic growth rate quantified by the parameter *r*_*p*_), while symbionts are obligate mutualists and can not grow in the absence of a host. We then extend this model to include interactions between an invasive host population (with biomass *p*_*i*_) and its invasive symbiont community (with biomass *m*_*i*_). We consider that native symbiont may exchange resources with native hosts, and invasive symbiont may exchange resources with native hosts. Competition between native and invasive hosts (*c*_*p*_ parameters) and between native and invasive symbionts (*c*_*m*_ parameters), may reduce their abundance, e.g., due to competition for host colonization between symbionts (Engelmoer et al., 2014; Smith et al., 2018), or due to host competition for light or other external resources (Craine and Dybzinski, 2013).

We obtain the following equations:

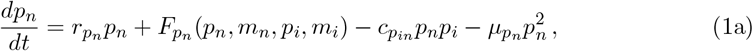

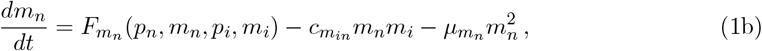

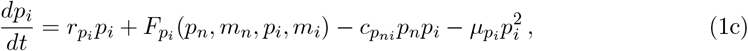

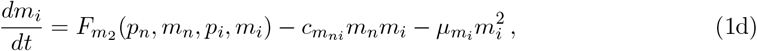

with functions 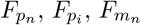, and 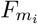defined as

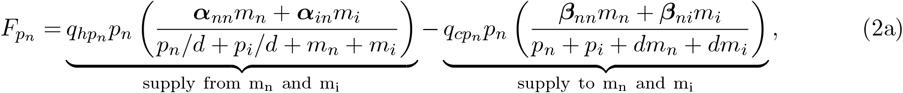

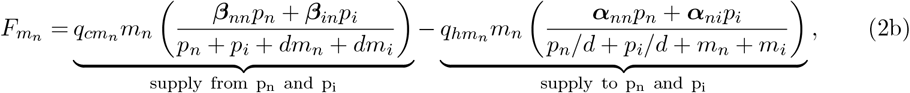

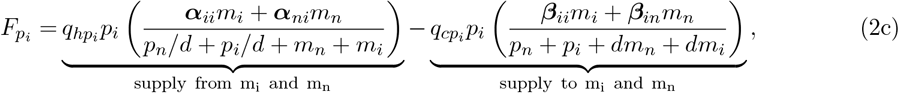

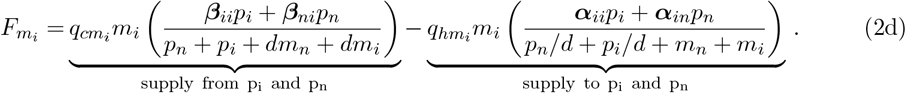

A schematic representation of this model is provided in Fig. 1(a). A table with a brief description of model parameters is provided in SI A.

**Fig. 1:**
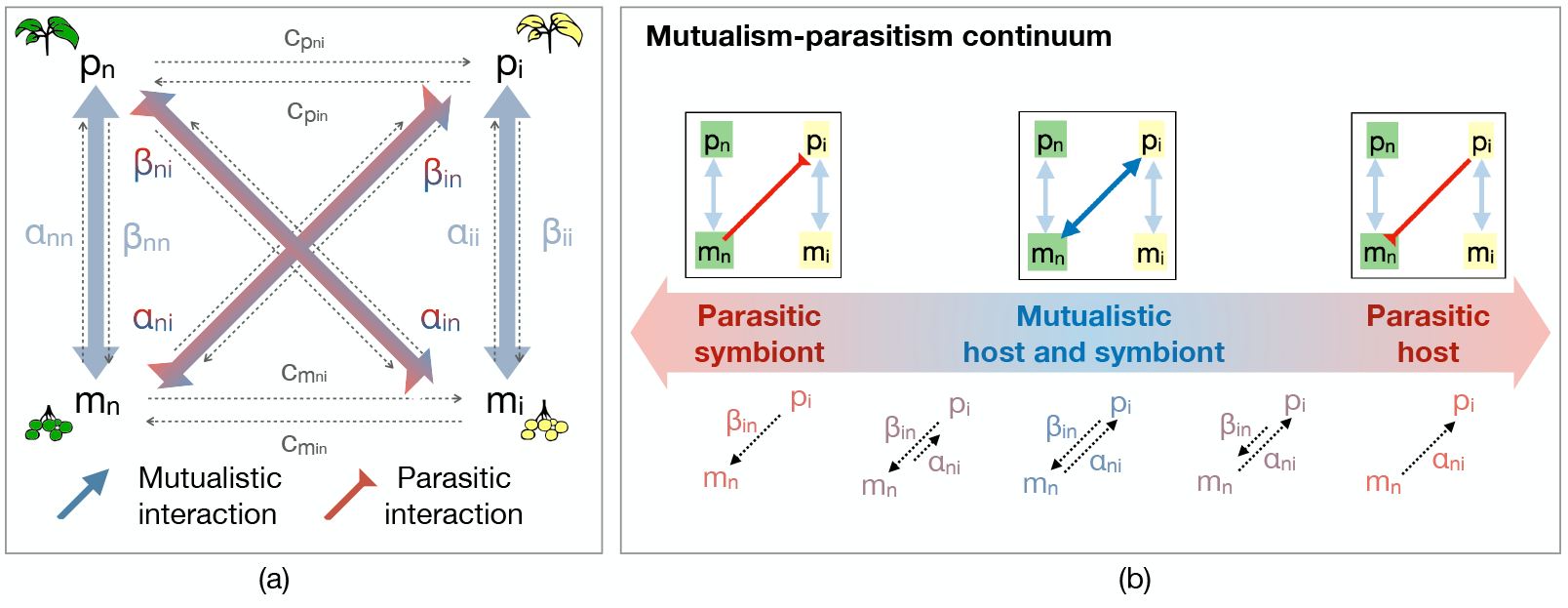
Schematic representation of the model of Eq. (1). A native microbial community *m*_*n*_ is associated with a host population *p*_*n*_. Resource exchange between hosts and symbionts is quantified by parameters *α*_*nn*_ (symbionts to hosts) and *β*_*nn*_ (hosts to symbionts). Similarly, a population of invasive hosts *p*_*i*_ exchanges resources with its associated invasive symbiont community *m*_*i*_ (parameters *α*_*ii*_ and *β*_*ii*_). Depending on the scenario considered, invasive hosts can also exchange resources with native symbionts, and so do invasive symbionts with native hosts (parameters *α*_*in*_, *α*_*ni*_, *β*_*in*_ and *β*_*ni*_). Blue and red arrows indicate whether host-symbiont interactions are parasitic or mutualistic for hosts and symbionts, as shown in (b). Additionally, native and invasive hosts compete with each other, with competition strength quantified by parameters 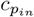 and 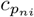, and so do native and invasive symbionts (parameters 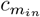 and 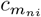).

The transfer of resources from symbionts to hosts and from hosts to symbionts is quantified by the *α*_*jk*_ and *β*_*jk*_ parameters respectively, with subindex *j* representing the supplying species (*n* for native, or *i* for invasive), and subindex *k* representing the receiving species. These parameters may represent particular traits in the receiving and supplying species, that quantify the resource exchange capacity of each one. For instance, microbes that provide lots of phosphorus to host plants and take lots of carbon from host plants are represented by large *α*_*j*_ and *β*_*j*_ parameters. Additionally, native and invasive species can differ in their intrinsic growth rate (parameters 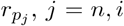), in their efficiency at converting the resource received or supplied into biomass (parameters 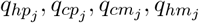, with *j* = *n, i*), and in the rate at which resources need to be diverted into maintenance of the existing biomass (parameters 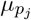 and 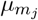, with *j* = *n, i*).

To explore the effect of host-symbiont association on invasion dynamics, we consider that parameters *α*_*in*_, *α*_*ni*_, *β*_*in*_ and *β*_*ni*_ can be zero or positive, depending on whether or not resource exchange between invasive symbionts/hosts and native hosts/symbionts is occurring.

If invasive hosts exchange nutrients with native symbionts, parameters *α*_*ni*_ (quantifying the resource supply from native symbionts to invasive hosts) and *β*_*in*_ (quantifying the resource supply from invasive hosts to native symbionts) will assume positive values. The relationship between how much a host receives from its associated symbionts (which depends on *α* parameters) and how much a host gives to its associated symbionts (which depends on *β* parameters) per unit time determines whether a host-symbiont relationship is mutualistic or parasitic for the host or the symbiont population (see Fig. 1b). Generally, the relationship is parasitic for the host population if *α* ≪ *β*, and parasitic for the symbiont community if *β* ≪ *α*, while the interaction is mutualistic for both hosts and symbionts, for *α* ≃ *β*. A detailed explanation of the quantitative criteria used to understand whether the exchange is mutualistic or parasitic is provided below in the ‘Model analysis and scenarios of interest’ section.

In addition to resource exchange parameters, resource supply also depends on host and symbiont densities, as determined by functions 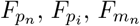 and 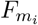, described in Eqs. (2a-d). Functions 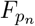and 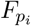 (Eqs. (2a,c)) tell us that when microbial biomass is much smaller than the total host biomass, the amount of resource that hosts can supply to each symbiont is limited by microbial biomass (i.e., by what symbiont can take), adjusted by the factor 1*/d*. Thus, each host species will supply to its symbionts an amount of resource proportional to symbiont biomass, and to the relative abundance of the host species in the whole host population, where the presence of other symbionts in the community may reduce this amount.

When host biomass is much smaller than the total symbiont biomass (adjusted by the factor 1*/d*) the amount of resource that a host can supply to its symbionts is limited by host biomass (e.g., by what the host can give). Each symbiont will supply to its host an amount of resource proportional to host biomass, and to the proportion of biomass occupied by the symbiont in the whole symbiont community. The presence of additional hosts may also reduce this amount. Analogously, functions 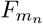 and 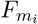(Eqs. (2b,d)) tell us that the amount of resource that symbionts supply to hosts is limited by symbiont biomass, when host biomass is large compared to the total symbiont biomass, and by symbiont biomass, when host biomass is smaller than the total symbiont biomass (adjusted by the factor *d*).

### 2.2 Model analysis and scenarios of interest

In our model, microbial contribution to host growth and host contribution to microbial growth vary along a continuum ranging from ‘parasitic’ to ‘mutualistic’ (see Fig. 1b). A certain host-symbiont association is defined as ‘mutualistic’ if the interaction increases the growth rate of both hosts and symbionts. An association is considered ‘parasitic’ if the growth rate of either hosts or symbionts, is decreased by the interaction.

Specifically, we consider the model of Eq. 1 in the case of a host population *p* (either *p*_*n*_ or *p*_*i*_) interacting with its microbial community *m* (either *m*_*n*_ or *m*_*i*_). We obtain:

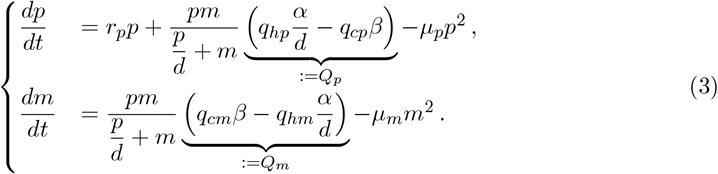

The association between *p* and *m* is mutualistic for *Q*_*p*_, *Q*_*m*_ *>* 0, parasitic for the host population if *Q*_*p*_ *<* 0 and *Q*_*m*_ *>* 0, and parasitic for the microbial community if *Q*_*p*_ *>* 0 and *Q*_*m*_ *<* 0. Note that the values of *Q*_*p*_ and *Q*_*m*_ depend on parameters *α* and *β*, which characterise the quality of the resource exchange between *p* and *m*.

We restrict our attention to the case where the association between native hosts and native symbionts (i.e., between *m*_*n*_ and *p*_*n*_) and between invasive hosts and invasive symbionts (i.e., between *m*_*i*_ and *p*_*i*_) is mutualistic. We then consider the possible outcomes when the associations between native symbionts *m*_*n*_ and invasive hosts *p*_*i*_ (Fig. 2, left) and between invasive symbionts *m*_*i*_ and native hosts *p*_*n*_ (Fig. 2, right) can be mutualistic or parasitic.

**Fig. 2:**
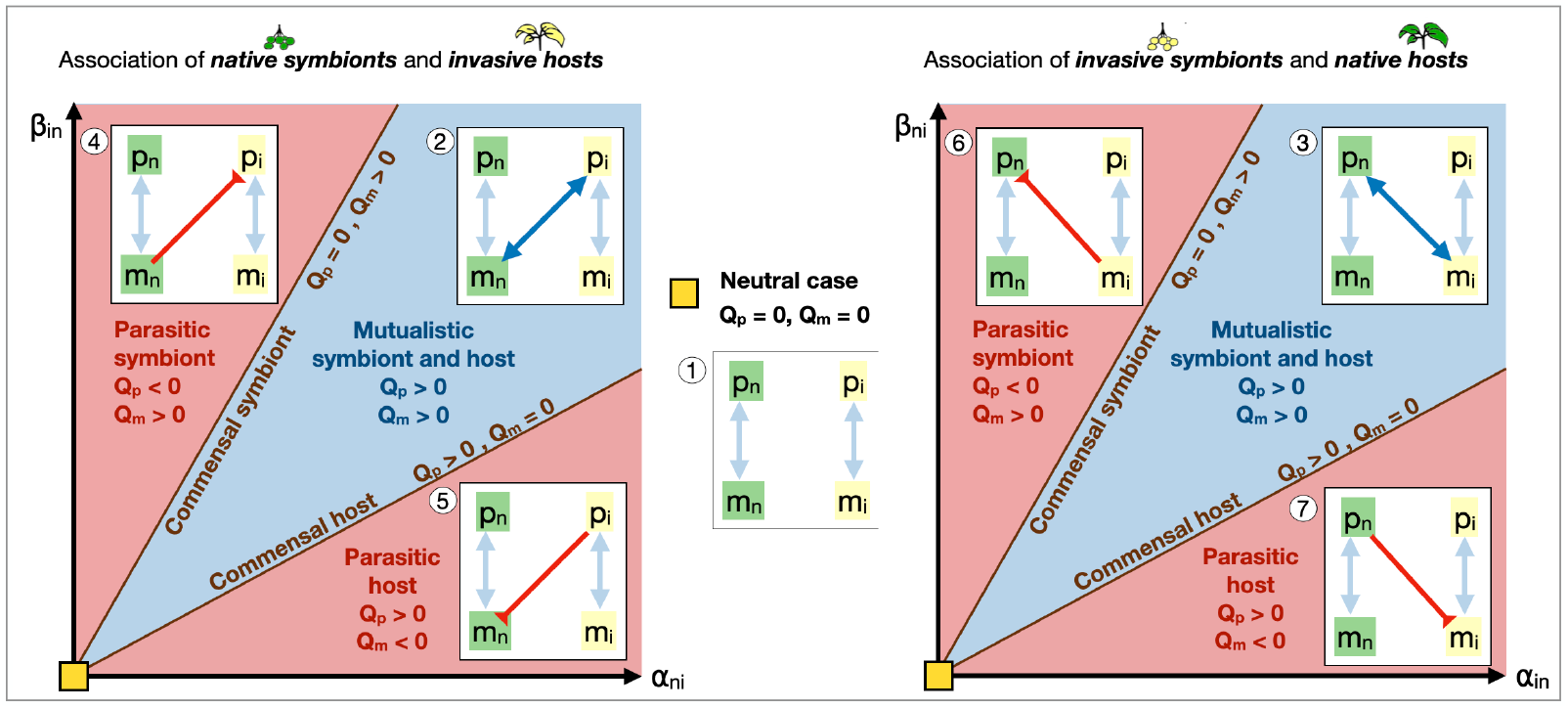
Novel host-symbiont interactions between native microbes and invasive hosts (left plot) or between native hosts and invasive microbes (right plot) can be parasitic or mutualistic, depending on the value of parameters *α*_*in*_, *α*_*ni*_, *β*_*ni*_ and *β*_*in*_, quantifying the resource exchange capacity of hosts and symbionts (see also Fig. 1). We characterize 7 different parameter regions, corresponding to scenarios 1-7, discussed in the manuscript. Left plot: Scenarios 2, 4, and 6; Middle plot: Scenario 1; Right plot: Scenarios 3, 6, and 7. The mathematical definition of the thresholds *Q*_*p*_ and *Q*_*m*_ are provided in Eq. (3) and Eqs. (4) and (5).

For instance, the association between a native host *p*_*n*_ and invasive microbial symbionts *m*_*i*_ is mutualistic as long as

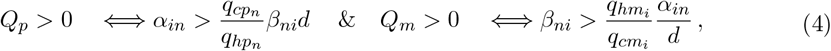

and parasitic for either the host or the symbiont if *Q*_*p*_ *<* 0 or *Q*_*m*_ *<* 0. Analogously, association between invasive hosts *p*_*i*_ and native microbial symbionts *m*_*n*_ is mutualistic for both hosts and symbionts if

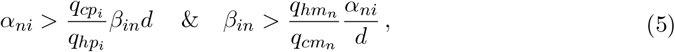

(see also SI B.2 for more details).

The straight lines defined by Eqs. (4) and (5), in the *α, β* parameter space, mark the borders between the different regions (parasitic symbiont, mutualistic symbiont and host, and parasitic host) in Fig. 2, in which interactions between an invasive host population *p*_*i*_ and a native symbiont community *m*_*n*_ is considered (left plot), or interactions between a native host population *p*_*n*_ and an invasive symbiont community *m*_*i*_ is considered (right plot). Equality in either terms of Eqs. (4) and (5) represents cases in which either hosts or symbionts are commensals.

We obtain seven different scenarios, which are identified in Fig. 2: ➀Invasive and native hosts do not share symbionts; a mutualistic association is observed between ➁ invasive hosts and a native symbionts, or ➂ native hosts and invasive symbionts; a parasitic association is observed, in which native/invasive symbionts exploit invasive/native hosts (➃ and ➅), or invasive/native hosts exploit native/invasive symbionts (➄ and ➆). The mathematical analysis of these scenarios and their biological significance will be discussed in section 3.2. Combinations of these scenarios will also be considered in section 3.3.

## 3 Results

### 3.1 Overview of model analysis

In the following, we discuss the interesting dynamics emerging from scenarios ➀ - ➆, which shed light on the several ways in which host-symbiont interactions can lead to invasion of hosts, symbionts, or both. Note that the directionality of the interactions is important, i.e., we distinguish between the effect of hosts on symbionts, and the effect of symbionts on hosts. The outcome may be the same in separate scenarios, but the mechanism leading to these outcomes differ.

A detailed mathematical analysis of these scenarios is presented in the SI B-E, which include: the analysis of the one host - one symbiont system, which is crucial to define the parasitic/mutualistic interactions between a host population and its symbionts community (section B); the analysis of the one host - two symbiont system (section C), which is important to understand the dynamics of a host population (native or invasive) associating with both native and invasive symbionts; the analysis of the one symbiont - two hosts system (section D), providing insights on the dynamics of a symbiont community (native or invasive) associating with native and invasive hosts; and the analysis of the two hosts - two symbionts system exposed according to the scenarios described in ➀ - ➆ (section E). We refer to invasion as a situation in which the successful establishment and persistance of invasive hosts, symbionts or both is possible. Exploration of alternative scenarios not presented in this paper, such as specific scenarios corresponding to different parameter combinations, can be conducted through numerical simulations of Eq. (1). For precisely this purpose, a user-friendly version of the code is made publicly available on the modelRxiv platform (Harris et al., 2022) (https://modelrxiv.org/model/YfndNX).

### 3.2 Scenarios of interest

#### ➀ Native and invasive species do not share symbionts nor hosts

When neither symbionts nor hosts are shared between native and invasive species, and competition within hosts and between symbionts is strong, competitive exclusion of one host-symbiont community, either the native or the invasive one, occurs through selection due to trait differences (represented in the model through differences in model parameters *α, β, q, c, μ*, or *r*_*p*_, which can be explored numerically), or through differences in initial abundance. When selection due to trait differences occurs, the host-symbiont association that provides the highest fitness to either hosts or symbionts outcompetes the other. Note that in this case, more mutualistic associations are expected to provide highest fitness, and therefore provide resilience against invasion. Differences in initial abundance can also favor one host-symbiont community over the other (see also Fig. E.1 ➀). Indeed, differences in the initial abundance of symbionts (or hosts) affect host (or symbiont) growth rate and, in turn, symbiont (or host) growth rate, providing a competitive advantage to the community with the largest abundance of hosts or symbionts.

#### ➁The association between native symbionts and invasive hosts is mutualistic

- *Association with native symbionts provides advantage to invasive hosts:* Association with native symbionts can increase the fitness of invasive hosts, and provide them with a competitive advantage that can lead to competitive exclusion of native hosts (Fig. 3b, right pathways, and SI E and Fig. E.1 ➁ for mathematical insights). Subsequently, invasive symbionts may outcompete native symbionts, e.g., if native symbionts are weakened by the absence of native hosts or if invasive symbionts are empowered by an increase in invasive hosts (Fig. 3a, far right pathway, leading to association between *p*_*i*_ and *m*_*i*_). Alternatively, invasive symbionts may be outcompeted by native symbionts (Fig. 3a, centre right pathway, leading to association between *p*_*i*_ and *m*_*n*_). A variation of this scenario in which invasive hosts are low quality mutualists is discussed in ➄ and in Fig. 4a. In this case, native symbionts receive a lower benefit from invasive hosts, compared to the benefit received from native hosts, but provide the same benefit to native and invasive hosts. Thus, introduced invasive hosts may grow rapidly by benefiting from the presence of a native microbial community and, indirectly, from the presence of native hosts. On the other hand, native symbionts and native hosts may suffer from the presence of invasive hosts, that benefit from native symbionts providing little in return. If this situation leads to a displacement of native hosts and symbionts, invasive hosts will no longer be able to acquire resources at little cost, and the remaining invasive host-symbiont community will have lower biomass than the initial native one. Note that when considering the case in which native and invasive species do not share symbionts (i.e., scenario ➀) less mutualistic host-symbiont associations are expected to have lower fitness with respect to more mutualistic host-symbiont associations, and are thus not likely to invade. When accounting for the formation of novel host-symbiont associations, however, co-invasion can be driven by the exploitation of existing host-symbiont associations, where invasive hosts indirectly exploit native hosts, by receiving resources from native symbionts at low cost.
- *Association with invasive hosts provides advantage to native symbionts:* Mutualistic association with invasive hosts can also provide a competitive advantage to native symbionts, that then outcompete invasive symbionts (Fig. 3b, left pathway). No invasion occurs if invasive hosts suffer from the disruption of invasive host-symbiont associations which causes them to be outcompeted by native hosts (Fig. 3b, far left pathway, leading to association between *p*_*n*_ and *m*_*n*_). However, if invasive symbionts are low quality mutualists to their own invasive hosts, their exclusion can provide a competitive advantage to invasive hosts, and cause the subsequent exclusion of native hosts. In this case, we observe the formation of novel associations between invasive hosts and native symbionts (Fig. 3b, center left pathway, leading to association between *p*_*i*_ and *m*_*n*_). Note that this scenario of microbial invasion through host replacement may be more likely to occur, as it can be observed through two different pathways, namely, the centre left and right pathways in Fig. 3b (see also SI E and Fig. E.1 ➁ for details on the dynamics).

**Fig. 3:**
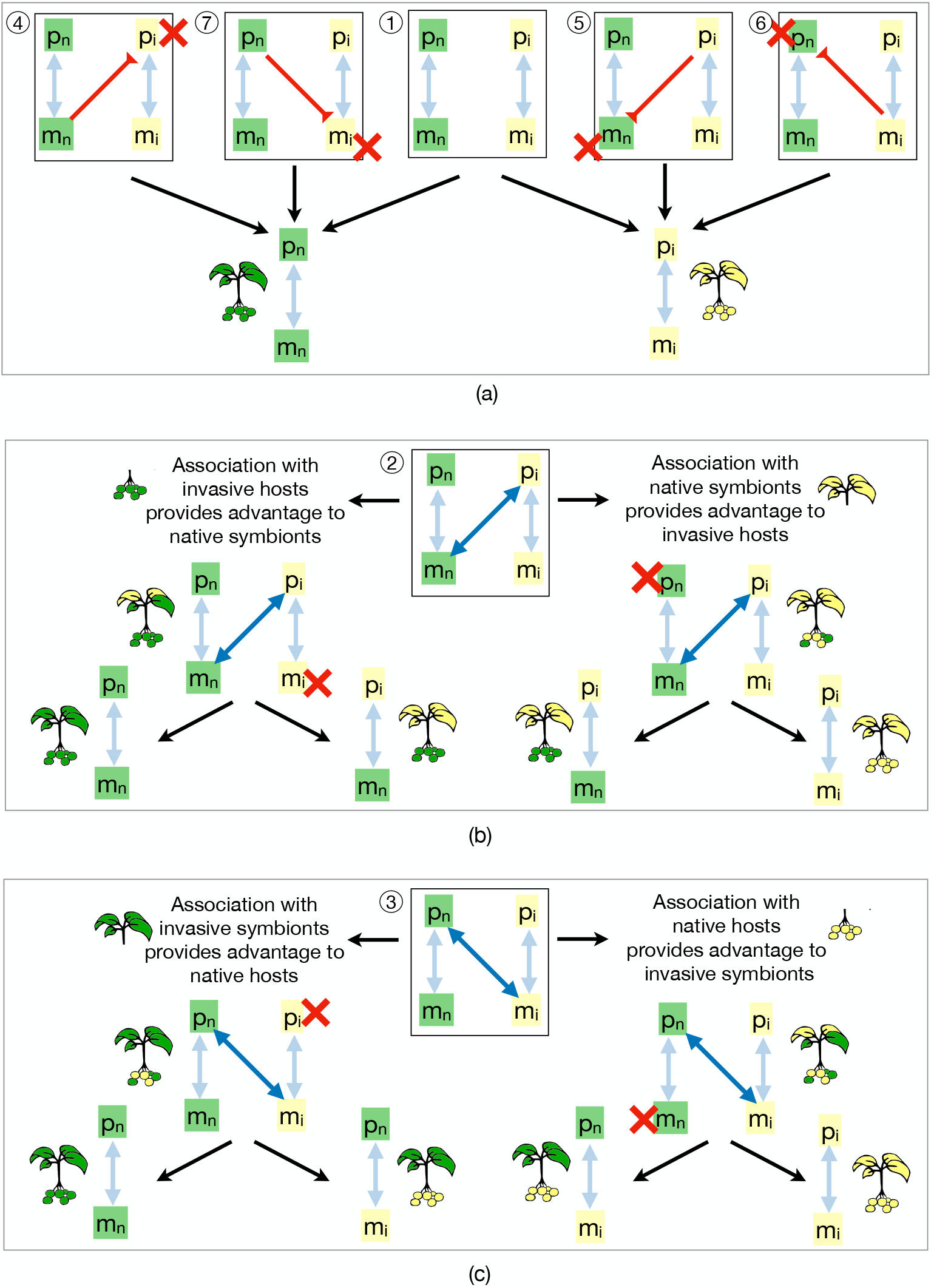
Possible dynamics of invasion occurring when native species form novel host-symbiont associations with invasive species, for each of the scenarios described in Fig. 2. (a) Co-invasion can be facilitated (scenarios 5 and 6) or prevented (scenarios 4 and 7) by the formation of parasitic associations between native and invasive species. Association of (b) native symbionts with invasive hosts (scenario 2) or of (c) invasive symbionts with native hosts (scenario 3) provides a competitive advantage to hosts or symbionts, where different outcomes are observed depending on whether hosts or microbes competitively exclude each other first.

**Fig. 4:**
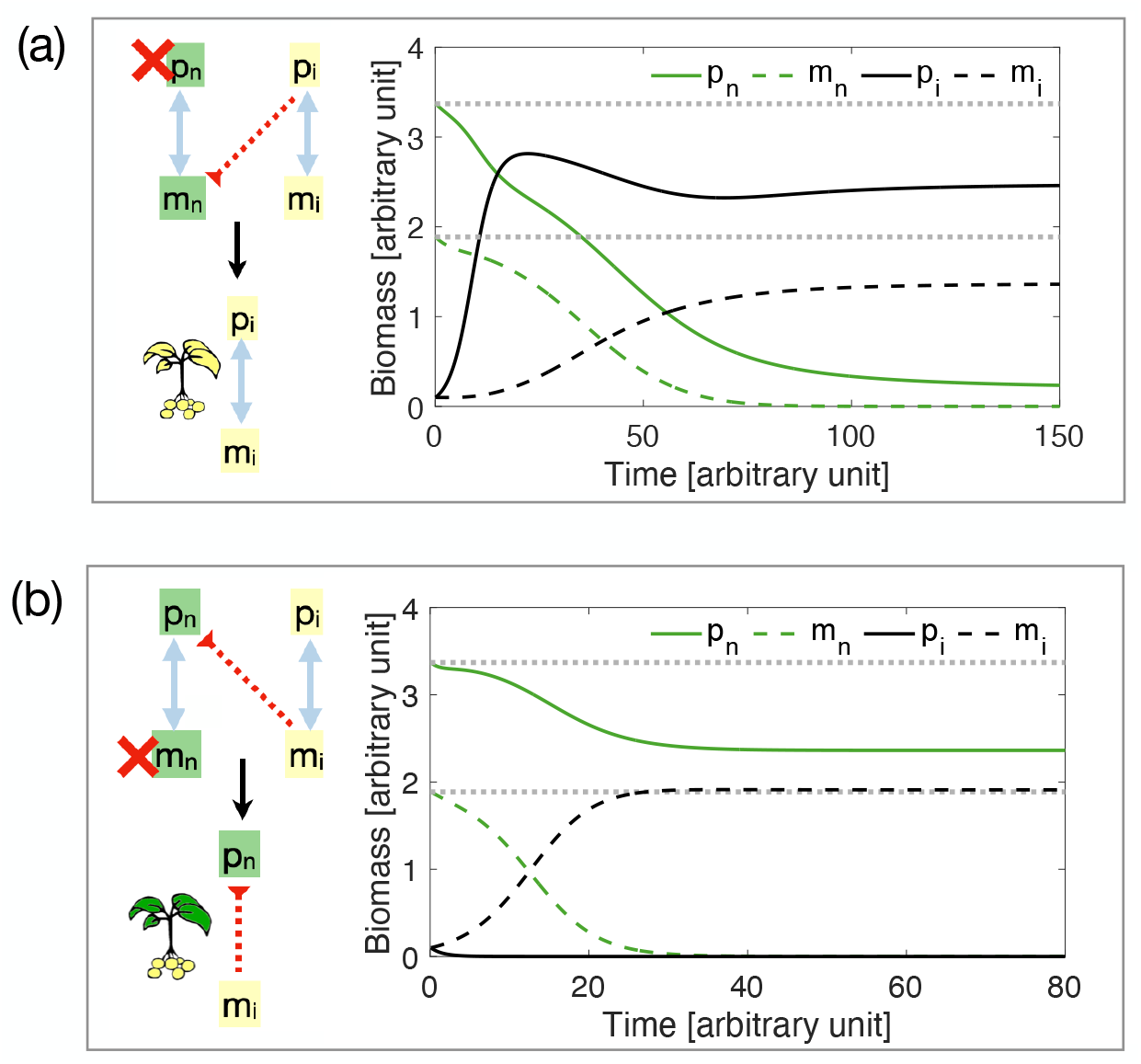
(a) Possible dynamics of co-invasion occurring when invasive hosts are low quality mutualists, and exploit native symbionts at the indirect expense of native hosts. In this case, the resulting community will have lower biomass with respect to the displaced invasive community (compare black curves and dotted grey horizontal lines). (b) Possible dynamics of microbial invasion occurring when native hosts associate with invasive symbionts that are low quality mutualists. In this case, invasive symbionts persist in the environment with native hosts, where the resulting community has lower biomass than the original one. Interactive simulation of these and related scenarios can be done through the modelRxiv platform (https://modelrxiv.org/model/YfndNX).

#### ➂The association between native hosts and invasive symbionts is mutualistic

- *Association with native hosts provides advantage to invasive symbionts:* If association with native hosts provides a competitive advantage to invasive symbionts, these may then then outcompete native symbionts (Fig. 3c, right pathways, and SI E and Fig. E.1 ➂ for details). The disruption of the association between native hosts and their symbionts may weaken native hosts, and lead to the establishment of invasive host-symbiont associations (Fig. 3c, far right pathway, leading to association between *p*_*i*_ and *m*_*i*_). Alternatively, microbial invasion may be observed if invasive hosts are outcompeted by native hosts, but invasive symbionts persist in association with native hosts (Fig. 3b center right pathway, leading to association between *p*_*n*_ and *m*_*i*_). A variation of the latter scenario is the situation in which invasive symbionts are not completely harmful to native hosts, but are low quality mutualists. In this case, the substitution of native symbionts with invasive symbionts may lead to a loss in the biomass of native hosts (Fig. 4b). Invasive microbes may then continue to be present in the environment and negatively affect ecosystem functionality long after the disappearance of their invasive hosts.
- *Association with invasive symbionts provides advantage to native hosts:* Association with invasive symbionts can also provide an advantage to native hosts, which then outcompete invasive hosts (Fig. 3c, left pathway and Fig. E.1 ➂). If invasive symbionts are strong competitors, they may subsequently exclude native symbionts, which would lead to the formation of novel associations between invasive symbionts and native hosts, and to microbial invasion (Fig. 3c center left pathway, leading to association between *p*_*n*_ and *m*_*i*_). This scenario may be more likely to occur, as it can be observed through two different pathways. No invasion occurs if invasive symbionts suffer from the absence of invasive hosts and are outcompeted by native symbionts (Fig. 3b far left pathway, leading to associations between *p*_*n*_ and *m*_*n*_).

#### ➃Native symbionts are parasitic to invasive hosts

The association of native symbionts with invasive hosts can provide biotic resistance to native host-symbiont associations (Fig. 3a, pathway ➃ and SI E and Fig. E.1 ➃). This can occur, for example, if parasites that are only slightly harmful to native hosts, e.g., because they have co-evolved with them, cause a strong reduction in the fitness, or death, of invasive hosts. The death of invasive hosts would then be followed by the death of their symbionts. Poor adaptation of invasive species to native symbionts can also lead to exploitation of invasive hosts by native symbionts. This can occur, for example, if native symbionts weaken invasive hosts by receiving a certain resource at low cost (see also Fig. D.2 for details).

#### ➄Invasive hosts are parasitic to native symbionts

If the invasive hosts harm microbial communities directly, e.g., by acquiring resources provided by native symbionts without giving anything in return, the interaction of invasive host and native symbionts would weaken native symbionts, and cause their displacement and the subsequent displacement of native hosts (Fig. 3a, pathway ➄ and SI E and Fig. E.1 ➄). This could be the case if the signalling dynamics between the host and the symbiont had not co-evolved. Thus, the host might be able to benefit from the symbiont against the latter’s best evolutionary interest, and due to the lack of previous interactions between the organisms, the symbiont might not be able to detect and respond adaptively to the situation, creating a new type of eco-evolutionary trap scenario (Ferriere and Legendre, 2013). If invasive hosts are not completely harmful to native symbionts, but are low quality mutualists, then their association with native symbionts can lead to exploitation of pre-existing native host-symbiont associations to invade, as discussed in ➁ and Fig. 4(a) (see also Fig. D.2 for details).

#### ➅Invasive symbionts are parasitic to native hosts

Co-invasion may be facilitated if invasive symbionts are harmful to native species, which would weaken native hosts and cause their competitive exclusion and the consequent exclusion of their symbionts (see Fig. 3a, pathway ➅, and SI E and Fig. E.1 ➅). This situation can occur, for example, if pathogens causing disease in native hosts are co-introduced with invasive hosts. A similar dynamics can occur if invasive symbionts are low quality mutualists, and weaken native causing their competitive exclusion by invasive hosts (see Fig. C.1).

#### ➆Native hosts are parasitic to invasive symbionts

Interaction of native hosts with invasive symbionts can provide resilience against invasion, if native hosts are harmful to invasive symbionts (Fig. 3a, pathway ➆ and SI E and Fig. E.1 ➆). Here, we consider that a native host can take advantage of invasive symbiont to the symbiont’s detriment, preventing the establishment of invasive symbionts and their hosts (see also Fig. C.1).

### 3.3 Combined scenarios

In the previous section, we discussed the possible effect of novel host-symbiont associations on invasion dynamics. Although these scenarios were considered in isolation, and we separately explored the effect of native species on invasive ones, and of invasive species on native ones, it is also possible that both native and invasive species form novel associations at the same time (see Fig. 5). For example, the acquisition of mutualistic native symbionts by invasive hosts (strengthening invasive hosts), and the transmission of parasitic symbionts from invasive to native hosts (weakening native hosts) can both occur (see Fig. 5, third row and third column, counting from the top left), leading to the combined effect of increasing the competitive ability of invasive hosts and decreasing the competitive ability of native hosts, facilitating host invasion. Similarly, a dynamic of microbial invasion is more likely to be observed if invasive symbionts are strengthened through the formation of novel association with native hosts, while native symbionts are harmed by invasive hosts (Fig. 5, second row, fourth column, counting from the top left).

**Fig. 5:**
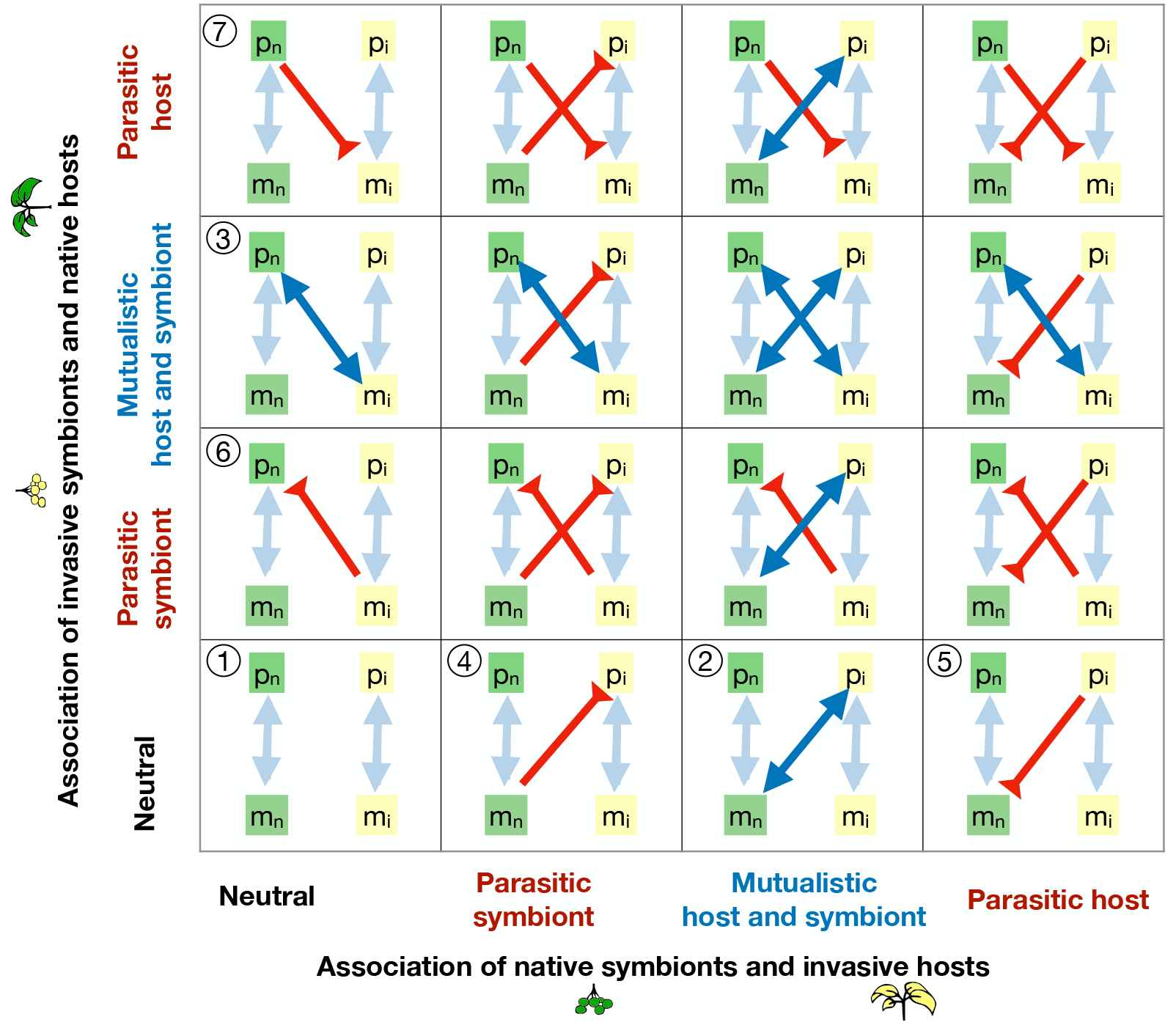
Schematic representation of possible interaction scenarios that can be explored within our framework. The seven numbered circles indicate the seven scenarios discussed in the results section. Other scenarios can be understood as combination of these seven. Interactive simulation of these and related scenarios, with our default parameters or user-defined values, can be done through the modelRxiv platform (https://modelrxiv.org/model/YfndNX).

## 4 Discussion

It has been known for some time that the formation of novel host-symbiont associations can facilitate invasion, as well as provide resilience to native communities (Bever et al., 2010; Dickie et al., 2017). We show here that these general observation can be dissected into several different mechanisms, which we use to create a global framework for broadening our understanding of invasion dynamics.

### 4.1 Host-symbiont interactions increase invasion risk

A growing number of empirical studies, particularly on plant-fungal associations, have shown that invasion can occur when invasive hosts form novel mutualistic associations with native symbionts, eventually increasing host competitive ability (Callaway et al., 2004; Tedersoo et al., 2007; Moeller et al., 2015; Shipunov et al., 2008). Strong evidence also shows that pathogens co-introduced with invasive hosts may weaken native host populations, which favours their competitive exclusion by invading hosts in a dynamic often referred to as ‘disease-mediated invasion’ (Anderson et al., 2004; Parker and Gilbert, 2004; Desprez-Loustau et al., 2007; Santini et al., 2013; Carnegie et al., 2016). We highlight the possibility that some of these mechanisms may occur not only when shared symbionts are either pathogens or mutualists, but also when the mutualistic quality of introduced symbionts differs from that of native symbionts. For instance, introduced symbionts that are slightly less mutualistic than those found in native communities might may lead to a decrease in biomass of native hosts (Bever, 2002) (e.g., see our Fig. 4b). Similarly, the acquisition of native symbionts that are slightly more mutualistic than the original invasive community can help a host to invade (see Fig. 4a). These changes in population growth and abundance may not be sufficient to directly drive a community to extinction (as for the acquisition of pathogenic microbes), but they may be enough to provide a competitive advantage to a population with respect to another and change the invasion dynamics (Levine et al., 2004).

In addition to highlighting the possible dynamics along the mutualism-parasitism continuum, we would like to emphasize the importance of considering invasion dynamics arising at the community level, that accounts for host-symbiont interactions, as well as host-host and symbiont-symbiont interactions. Considering only host-symbiont interactions may lead to the misconception that only the host-symbiont community that provides the highest fitness to their hosts and symbionts may co-invade and displace native host-symbiont communities, as observed in some instances (Dickie et al., 2010; Nunez and Dickie, 2014; Hayward et al., 2015). However, when accounting for the possibility that symbiont disruption and exchange among native and invasive species may occur (Dickie et al., 2017; Catford et al., 2009; Mitchell et al., 2006), co-invasion of a less fit host-symbiont community may be observed (Fig. 4a). Thus, understanding the whole range of possible outcomes following the introduction of a new species requires to embrace a community perspective that accounts for interactions among multiple hosts and symbionts (Dickie et al., 2017; Fahey and Flory, 2022).

In some instances, the formation of novel host-symbiont interactions may lead to changes in total community biomass with long-term repercussions on ecosystem functionality (Nunez and Dickie, 2014; Lovett et al., 2006; Dickie et al., 2011; Mitchell and Power, 2003; Cobb and Rizzo, 2016; Preston et al., 2016; Ehrenfeld, 2010). Although this reduction in community biomass is well-known for pathogen spread (Lovett et al., 2006; Mitchell and Power, 2003; Cobb and Rizzo, 2016; Preston et al., 2016), here we present alternative mechanisms that can lead to an invasion-driven biomass decrease through higher-order interactions (Billick and Case, 1994; Mayfield and Stouffer, 2017). For example, if invasive hosts provide a reduced reward as compared to natives (Hoffman and Mitchell, 1986; Mummey and Rillig, 2006; Hausmann and Hawkes, 2009; Vogelsang and Bever, 2009), associations between invasive hosts and native symbionts will lead to the direct exploitation of the resources provided by native symbionts. In addition, there will be indirect exploitation of more mutualistic native hosts that invested resources in the growth of a large native microbial community from which it can no longer fully benefit. Thus, in this case, an increase in the abundance of invasive hosts, facilitated by the presence of native host-symbiont communities, occurs in conjunction with a decrease in the abundance of native symbionts and their hosts (e.g., see Fig. 4a).

### 4.2 Host-symbiont interactions increase community resilience

Less studied than the role of host-symbiont associations in invasion dynamics is their role in providing resilience to native communities (Van der Putten et al., 2010; Zenni and Nuñez, 2013; Levine et al., 2004). In spite of its importance to community assembly (Wu et al., 2024), invasion failure remains poorly understood in practice Diez et al. (2009); Zenni and Nuñez (2013). Given that novel host-symbiont associations may be the key to understand invasion success, they may also underlie mechanisms providing resistance to invasion. Indeed, mechanisms of symbiont disruption and replacement, as well as differences in community composition and emerging properties following species introduction, may lead to changes in resource exchange dynamics between hosts and symbionts and, possibly, provide resistance to invasion (Levine et al., 2004; Dinoor and Eshed, 1984; Beckstead and Parker, 2003; Knevel et al., 2004).

A clear example of host-symbiont associations driving invasion failure is the transmission of native pathogens to invasive plants (Hood et al., 2008; Piou et al., 2002). This novel association can provide biotic resistance against invaders (Mordecai, 2013; Flory and Clay, 2013; Prevéy and Seastedt, 2015). Other, more complex, dynamics of biotic resistance occur when native symbionts, which have co-evolved with native hosts, are highly mutualistic to native hosts but not to invasive hosts (Bunn et al., 2015; Moora et al., 2011). Here we show that association with these low-quality mutualistic symbionts can harm invasive hosts, and allow for their competitive exclusion by native hosts (as described in 2 and 4).

Finally, less competitive host-symbiont pairs may resist invasion by associating with mutualistic invasive symbionts (Mordecai, 2013; Flory and Clay, 2013). Reports of spread of invasive symbionts in native habitats are numerous (Dickie et al., 2016; Wolfe and Pringle, 2012; Berch et al., 2017; Mallon et al., 2015; Hart et al., 2017; Golan et al., 2024). However, the ecological consequences of these new associations are unclear. Our framework can guide new empirical research to understand under which circumstances these associations can provide natives with biotic resistance to host invasion, and when they may instead lead to a risk of facilitating symbiont invasion.

### 4.3 Framework limitations and possible extensions

Our framework allows for the mechanistic exploration of the interactions between microbes and their hosts, and can produce a rich variety of theoretical scenarios. Parameterization based on realistic biological scenarios that consider the specificity of host-symbiont interactions and variability in their contribution to fitness can provide insights into possible invasion dynamics through numerical simulations (e.g., through the tool we provide on the modelRxiv platform Harris et al. (2022)). These simulations, can help us determine which scenarios are more likely to be observed in particular settings.

The framework presented here also provides a strong basis for new mathematical investigation. Our model accounts for multiple levels of interactions -such as interactions between hosts, between symbionts, and between hosts and symbionts-, it considers biologically plausible density-dependent functional responses (as shown in Eqs. (2)), as well as an asymmetry in their dependence on the mutualism (i.e., obligate or facultative). Thus, the effect of higher-order interactions in our system may differ from what observed in previous studies that considered only microbial or only host communities, and that used linear, instead of density-dependent, functional responses (e.g., Gibbs et al. (2022)). Additionally, an extension of this framework to account for associations between multiple hosts and symbionts (e.g., to consider a microbial community composed of multiple microbial strains in which invasive hosts preferentially support certain microbial strains over others (Callaway et al., 2001; Bever, 2002; Kohout et al., 2011)), could help us investigate the effect of higher-order interactions on coexistence and diversity of large host-symbiont communities. Such an extension to our framework is presented in SI F) Other possible extensions include the evolution of host-symbiont associations, such as the evolution of host adaptation to pathogens (Thrall et al., 2002), or ‘parasite-spillback’ mechanisms (Flory and Clay, 2013; Strauss et al., 2012; Kelly et al., 2009; Mangla et al., 2008; Day et al., 2016), where invasive hosts associate with native pathogens that increase in abundance on invasive hosts, leading to increased colonization of native hosts.

## 5 Conclusion

Multi-species interactions involving native and invasive hosts and their symbionts can play an important role in invasion dynamics, and our ability to accurately evaluate invasion risks has been shown to depend their intricate dynamics. While classical theory has mostly focus on the individual invasion of either host species or their microbial symbionts, consideration on the possible formation of novel host-symbiont associations between native and non-native symbionts and their native and non-native hosts can shed light on new modalities of invasion that have not been considered so far. Thus, less mutualistic hosts and symbionts can invade through exploitation of pre-existing symbiotic relationships, or introduced microbes can increase their invasiveness through association with native hosts. We present a mechanistic mathematical framework that explicitly accounts for host-symbiont mutualistic and parasitic associations, to deepen our understanding of invasion biology in ecological communities and guide empirical research.

## Statements and declarations

- The authors have no competing interests to declare that are relevant to the content of this article.
- All authors certify that they have no affiliations with or involvement in any organization or entity with any financial interest or non-financial interest in the subject matter or materials discussed in this manuscript.
- The authors have no financial or proprietary interests in any material discussed in this article.
- This study did not require any human involvement, and no human data was taken during the study
- Data availability statement: The datasets generated or analyzed during the study are available in this published article. All the code developed for the manuscript is available on GitHub. The code is novel and original, and has been developed by author Maria M. Martignoni (2023) and can be found at https://github.com/nanomaria/microbiallymediatedinvasion.
- Interactive reproduction and re-parametrization of results can also be done through the modelRxiv platform (https://modelrxiv.org/model/YfndNX).
- Fundings: MMM was funded by the Azrieli Foundation. OK was funded by the Israel Science Foundation (ISF) (grant number 1826/20), the Gordon and Betty Moore Foundation, the United States-Israel Binational Science Foundation (BSF), and The Minerva Center for the Study of Population Fragmentation. RCT was funded by the Natural Sciences and Engineering Research Council (NSERC) of Canada Discovery Grants Program, grant number RGPIN-2022-03589, and the University of British Columbia Okanagan Institute for Biodiversity, Resilience, and Ecosystem Services. JG was funded by ModEcoEvo project funded by the Université Savoie Mont-Blanc and the ANR project ReaCh (ANR-23-CE40-0023-01).
- Author’s contributions: MMM: Conceptualization, Methodology, Formal analysis, Investigation, Visualization, Writing - Original Draft, Writing - Review & Editing. JG: Conceptualization, Methodology, Formal analysis, Investigation, Visualization, Writing - Review & Editing. RT: Conceptualization, Methodology, Writing - Review & Editing. KH: Results visualization on modelRxiv. OK: Conceptualization, Methodology, Writing - Review & Editing. All authors agree with the present manuscript.

## References

Amsellem, L., Brouat, C., Duron, O., Porter, S. S., Vilcinskas, A., and Facon, B. Importance of microorganisms to macroorganisms invasions: is the essential invisible to the eye?(the little prince, a. de saint-exupéry, 1943). In Advances in ecological research, volume 57, pages 99–146. Elsevier, 2017.

Anderson, P. K., Cunningham, A. A., Patel, N. G., Morales, F. J., Epstein, P. R., and Daszak, P. Emerging infectious diseases of plants: pathogen pollution, climate change and agrotechnology drivers. Trends in Ecology & Evolution, 19(10):535–544, 2004.

Bascompte, J. and Jordano, P. Mutualistic networks. Princeton University Press, 2013.

Bascompte, J. and Olesen, J. M. Mutualistic networks. In Bronstein, J. L., editor, Mutualism. Oxford University Press, 07 2015. ISBN 9780199675654.

Bastolla, U., Fortuna, M. A., Pascual-García, A., Ferrera, A., Luque, B., and Bascompte, J. The architecture of mutualistic networks minimizes competition and increases biodiversity. Nature, 458(7241):1018–1020, 2009.

Beckstead, J. and Parker, I. M. Invasiveness of ammophila arenaria: release from soil-borne pathogens? Ecology, 84(11):2824–2831, 2003.

Berch, S. M., Kroeger, P., and Finston, T. The death cap mushroom (amanita phalloides) moves to a native tree in victoria, british columbia. Botany, 95(4):435–440, 2017.

Bever, J. D. Negative feedback within a mutualism: host–specific growth of mycorrhizal fungi reduces plant benefit. Proceedings of the Royal Society of London. Series B: Biological Sciences, 269(1509):2595–2601, 2002.

Bever, J. D., Westover, K. M., and Antonovics, J. Incorporating the soil community into plant population dynamics: the utility of the feedback approach. Journal of Ecology, pages 561–573, 1997.

Bever, J. D., Dickie, I. A., Facelli, E., Facelli, J. M., Klironomos, J., Moora, M., Rillig, M. C., Stock, W. D., Tibbett, M., and Zobel, M. Rooting theories of plant community ecology in microbial interactions. Trends in Ecology & Evolution, 25(8):468–478, 2010.

Billick, I. and Case, T. J. Higher order interactions in ecological communities: what are they and how can they be detected? italic>Ecology, 75(6):1529–1543, 1994.

Bunn, R. A., Ramsey, P. W., and Lekberg, Y. Do native and invasive plants di?er in their interactions with arbuscular mycorrhizal fungi? a meta-analysis. Journal of Ecology, 103 (6):1547–1556, 2015.

Callaway, R., Newingham, B., Zabinski, C. A., and Mahall, B. E. Compensatory growth and competitive ability of an invasive weed are enhanced by soil fungi and native neighbours. Ecology Letters, 4(5):429–433, 2001.

Callaway, R. M., Thelen, G. C., Rodriguez, A., and Holben, W. E. Soil biota and exotic plant invasion. Nature, 427(6976):731–733, 2004.

Campbell, C., Yang, S., Shea, K., and Albert, R. Topology of plant-pollinator networks that are vulnerable to collapse from species extinction. Physical Review E, 86(2):021924, 2012.

Carnegie, A. J., Kathuria, A., Pegg, G. S., Entwistle, P., Nagel, M., and Giblin, F. R. Impact of the invasive rust puccinia psidii (myrtle rust) on native myrtaceae in natural ecosystems in australia. Biological Invasions, 18:127–144, 2016.

Catford, J. A., Jansson, R., and Nilsson, C. Reducing redundancy in invasion ecology by integrating hypotheses into a single theoretical framework. Diversity and distributions, 15 (1):22–40, 2009.

Chiarello, M., Bucholz, J. R., McCauley, M., Vaughn, S. N., Hopper, G. W., Sánchez González, I., Atkinson, C. L., Lozier, J. D., and Jackson, C. R. Environment and co-occurring native mussel species, but not host genetics, impact the microbiome of a freshwater invasive species (corbicula fluminea). Frontiers in Microbiology, 13:800061, 2022.

Cobb, R. C. and Rizzo, D. M. Litter chemistry, community shift, and non-additive e?ects drive litter decomposition changes following invasion by a generalist pathogen. Ecosystems, 19:1478–1490, 2016.

Compant, S., Samad, A., Faist, H., and Sessitsch, A. A review on the plant microbiome: Ecology, functions, and emerging trends in microbial application. Journal of Advanced Research, 19:29–37, 2019.

Craine, J. M. and Dybzinski, R. Mechanisms of plant competition for nutrients, water and light. Functional Ecology, 27(4):833–840, 2013.

Dawkins, K. and Esiobu, N. Emerging insights on brazilian pepper tree (schinus terebinthi-folius) invasion: the potential role of soil microorganisms. Frontiers in Plant Science, page 712, 2016.

Day, N. J., Dunfield, K. E., and Antunes, P. M. Fungi from a non-native invasive plant increase its growth but have di?erent growth e?ects on native plants. Biological invasions, 18:231–243, 2016.

Desprez-Loustau, M.-L., Robin, C., Buee, M., Courtecuisse, R., Garbaye, J., Suffert, F., Sache, I., and Rizzo, D. M. The fungal dimension of biological invasions. Trends in Ecology & Evolution, 22(9):472–480, 2007.

Dickie, I. A., Bolstridge, N., Cooper, J. A., and Peltzer, D. A. Co-invasion by pinus and its mycorrhizal fungi. New Phytologist, 187(2):475–484, 2010.

Dickie, I. A., Yeates, G. W., St. John, M. G., Stevenson, B. A., Scott, J. T., Rillig, M. C., Peltzer, D. A., Orwin, K. H., Kirschbaum, M. U., Hunt, J. E., et al. Ecosystem service and biodiversity trade-o?s in two woody successions. Journal of Applied Ecology, 48(4): 926–934, 2011.

Dickie, I. A., Nuñez, M. A., Pringle, A., Lebel, T., Tourtellot, S. G., and Johnston, P. R. Towards management of invasive ectomycorrhizal fungi. Biological Invasions, 18:3383–3395, 2016.

Dickie, I. A., Bufford, J. L., Cobb, R. C., Desprez-Loustau, M.-L., Grelet, G., Hulme, P. E., Klironomos, J., Makiola, A., Nuñez, M. A., Pringle, A., et al. The emerging science of linked plant–fungal invasions. New Phytologist, 215(4):1314–1332, 2017.

Diez, J. M., Williams, P. A., Randall, R. P., Sullivan, J. J., Hulme, P. E., and Duncan, R. P. Learning from failures: testing broad taxonomic hypotheses about plant naturalization. Ecology letters, 12(11):1174–1183, 2009.

Díez, J. Invasion biology of australian ectomycorrhizal fungi introduced with eucalypt plantations into the iberian peninsula. Biological Invasions, 7:3–15, 2005.

Dinoor, A. and Eshed, N. The role and importance of pathogens in natural plant communities. Annual Review of Phytopathology, 22(1):443–466, 1984.

Dunn, A. M. and Hatcher, M. J. Parasites and biological invasions: parallels, interactions, and control. Trends in Parasitology, 31(5):189–199, 2015.

Ehrenfeld, J. G. Ecosystem consequences of biological invasions. Annual review of ecology, evolution, and systematics, 41:59–80, 2010.

Engelmoer, D. J., Behm, J. E., and Toby Kiers, E. Intense competition between arbuscular mycorrhizal mutualists in an in vitro root microbiome negatively affects total fungal abundance. Molecular Ecology, 23(6):1584–1593, 2014.

Fahey, C. and Flory, S. L. Soil microbes alter competition between native and invasive plants. Journal of Ecology, 110(2):404–414, 2022.

Ferriere, R. and Legendre, S. Eco-evolutionary feedbacks, adaptive dynamics and evolutionary rescue theory. Philosophical Transactions of the Royal Society B: Biological Sciences, 368 (1610):20120081, 2013.

Fitzpatrick, C. R., Salas-González, I., Conway, J. M., Finkel, O. M., Gilbert, S., Russ, D., Teixeira, P. J. P. L., and Dangl, J. L. The plant microbiome: from ecology to reductionism and beyond. Annual Review of Microbiology, 74:81–100, 2020.

Flory, S. L. and Clay, K. Pathogen accumulation and long-term dynamics of plant invasions. Journal of Ecology, 101(3):607–613, 2013.

Gibbs, T., Levin, S. A., and Levine, J. M. Coexistence in diverse communities with higher-order interactions. Proceedings of the National Academy of Sciences, 119(43):e2205063119, 2022.

Gibbs, T. L., Gellner, G., Levin, S. A., McCann, K. S., Hastings, A., and Levine, J. M. Can higher-order interactions resolve the species coexistence paradox? bioRxiv, pages 2023–06, 2023.

Goedknegt, M. A., Schuster, A.-K., Buschbaum, C., Gergs, R., Jung, A. S., Luttikhuizen, P. C., Van der Meer, J., Troost, K., Wegner, K. M., and Thieltges, D. W. Spillover but no spillback of two invasive parasitic copepods from invasive pacific oysters (crassostrea gigas) to native bivalve hosts. Biological Invasions, 19:365–379, 2017.

Golan, J., Wang, Y.-W., Adams, C. A., Cross, H., Elmore, H., Gardes, M., Gonçalves, S. C., Hess, J., Richard, F., Wolfe, B., et al. Death caps (amanita phalloides) frequently establish from sexual spores, but individuals can grow large and live for more than a decade in invaded forests. New Phytologist, 242(4):1753–1770, 2024.

Grünwald, N. J., Garbelotto, M., Goss, E. M., Heungens, K., and Prospero, S. Emergence of the sudden oak death pathogen phytophthora ramorum. Trends in Microbiology, 20(3): 131–138, 2012.

Gubbins, S., Gilligan, C. A., and Kleczkowski, A. Population dynamics of plant–parasite interactions: thresholds for invasion. Theoretical Population Biology, 57(3):219–233, 2000.

Harris, K. D., Hadari, G., and Greenbaum, G. modelrxiv: A platform for the distribution, computation and interactive display of models. bioRxiv, pages 2022–02, 2022.

Hart, M. M., Antunes, P. M., and Abbott, L. K. Unknown risks to soil biodiversity from commercial fungal inoculants. Nature ecology & evolution, 1(4):0115, 2017.

Hausmann, N. T. and Hawkes, C. V. Plant neighborhood control of arbuscular mycorrhizal community composition. New Phytologist, 183(4):1188–1200, 2009.

Hayward, J., Horton, T. R., and Nuñez, M. A. Ectomycorrhizal fungal communities coinvading with p inaceae host plants in a rgentina: G ringos bajo el bosque. New Phytologist, 208(2): 497–506, 2015.

Ho?man, M. and Mitchell, D. The root morphology of some legume spp. in the south-western cape and the relationship of vesicular-arbuscular mycorrhizas with dry mass and phosphorus content of acacia saligna seedlings. South African Journal of Botany, 52(4):316–320, 1986.

Holland, J. N. Population ecology of mutualism. In Bronstein, J. L., editor, Mutualism. Oxford University Press, 07 2015. ISBN 9780199675654.

Holland, J. N. and DeAngelis, D. L. A consumer–resource approach to the density-dependent population dynamics of mutualism. Ecology, 91(5):1286–1295, 2010.

Hood, I., Petrini, L., and Gardner, J. Colonisation of woody material in pinus radiata plantations by armillaria novae-zelandiae basidiospores. Australasian Plant Pathology, 37:347–352, 2008.

Hui, C., Richardson, D. M., Landi, P., Minoarivelo, H. O., Garnas, J., and Roy, H. E. Defining invasiveness and invasibility in ecological networks. Biological Invasions, 18:971–983, 2016.

Jack, C. N., Friesen, M. L., Hintze, A., and Sheneman, L. Third-party mutualists have contrasting e?ects on host invasion under the enemy-release and biotic-resistance hypotheses. Evolutionary Ecology, 31:829–845, 2017.

Kandlikar, G. S., Johnson, C. A., Yan, X., Kraft, N. J., and Levine, J. M. Winning and losing with microbes: how microbially mediated fitness di?erences influence plant diversity. Ecology letters, 22(8):1178–1191, 2019.

Kelly, D., Paterson, R., Townsend, C., Poulin, R., and Tompkins, D. Parasite spillback: a neglected concept in invasion ecology? Ecology, 90(8):2047–2056, 2009.

Knevel, I. C., Lans, T., Menting, F. B., Hertling, U. M., and van der Putten, W. H. Release from native root herbivores and biotic resistance by soil pathogens in a new habitat both a?ect the alien ammophila arenaria in south africa. Oecologia, 141:502–510, 2004.

Kohout, P., Sy`korová, Z., Bahram, M., Hadincová, V., Albrechtová, J., Tedersoo, L., and Vohník, M. Ericaceous dwarf shrubs affect ectomycorrhizal fungal community of the invasive pinus strobus and native pinus sylvestris in a pot experiment. Mycorrhiza, 21:403–412, 2011.

Levine, J. M., Adler, P. B., and Yelenik, S. G. A meta-analysis of biotic resistance to exotic plant invasions. Ecology letters, 7(10):975–989, 2004.

Lewis, M. A., Petrovskii, S. V., Potts, J. R., et al. The mathematics behind biological invasions, volume 44. Springer, 2016.

Lovett, G. M., Canham, C. D., Arthur, M. A., Weathers, K. C., and Fitzhugh, R. D. Forest ecosystem responses to exotic pests and pathogens in eastern north america. BioScience, 56(5):395–405, 2006.

Mallon, C. A., Van Elsas, J. D., and Salles, J. F. Microbial invasions: the process, patterns, and mechanisms. Trends in Microbiology, 23(11):719–729, 2015.

Mangla, S., Inderjit, and Callaway, R. M. Exotic invasive plant accumulates native soil pathogens which inhibit native plants. Journal of Ecology, 96(1):58–67, 2008.

Martignoni, M. M. and Kolodny, O. Microbiome transfer from native to invasive species may increase invasion risk and shorten invasion lag. bioRxiv, pages 2023–08, 2023.

Martignoni, M. M., Hart, M. M., Garnier, J., and Tyson, R. C. Parasitism within mutualist guilds explains the maintenance of diversity in multi-species mutualisms. Theoretical Ecology, 13:615–627, 2020.

Mayfield, M. M. and Stouffer, D. B. Higher-order interactions capture unexplained complexity in diverse communities. Nature Ecology & Evolution, 1(3):0062, 2017.

Médoc, V., Firmat, C., Sheath, D., Pegg, J., Andreou, D., and Britton, J. Parasites and biological invasions: predicting ecological alterations at levels from individual hosts to whole networks. In Advances in Ecological Research, volume 57, pages 1–54. Elsevier, 2017.

Minoarivelo, H. O. and Hui, C. Invading a mutualistic network: to be or not to be similar. Ecology and Evolution, 6(14):4981–4996, 2016a.

Minoarivelo, H. and Hui, C. Trait-mediated interaction leads to structural emergence in mutualistic networks. Evolutionary ecology, 30:105–121, 2016b.

Mitchell, C. E. and Power, A. G. Release of invasive plants from fungal and viral pathogens. Nature, 421(6923):625–627, 2003.

Mitchell, C. E., Agrawal, A. A., Bever, J. D., Gilbert, G. S., Hufbauer, R. A., Klironomos, J. N., Maron, J. L., Morris, W. F., Parker, I. M., Power, A. G., et al. Biotic interactions and plant invasions. Ecology letters, 9(6):726–740, 2006.

Moeller, H. V., Dickie, I. A., Peltzer, D. A., and Fukami, T. Mycorrhizal co-invasion and novel interactions depend on neighborhood context. Ecology, 96(9):2336–2347, 2015.

Moora, M., Berger, S., Davison, J., Ö pik, M., Bommarco, R., Bruelheide, H., Kühn, I., Kunin, W. E., Metsis, M., Rortais, A., et al. Alien plants associate with widespread generalist arbuscular mycorrhizal fungal taxa: evidence from a continental-scale study using massively parallel 454 sequencing. Journal of Biogeography, 38(7):1305–1317, 2011.

Mordecai, E. A. Despite spillover, a shared pathogen promotes native plant persistence in a cheatgrass-invaded grassland. Ecology, 94(12):2744–2753, 2013.

Mummey, D. L. and Rillig, M. C. The invasive plant species centaurea maculosa alters arbuscular mycorrhizal fungal communities in the field. Plant and Soil, 288:81–90, 2006.

Nunez, M. A. and Dickie, I. A. Invasive belowground mutualists of woody plants. Biological Invasions, 16:645–661, 2014.

Nuñstring-nameez, M. A., Horton, T. R., and Simberloff, D. Lack of belowground mutualisms hinders pinaceae invasions. Ecology, 90(9):2352–2359, 2009.

Panzavolta, T., Bracalini, M., Benigno, A., and Moricca, S. Alien invasive pathogens and pests harming trees, forests, and plantations: Pathways, global consequences and management. Forests, 12(10):1364, 2021.

Parepa, M., Schaffner, U., and Bossdorf, O. Help from under ground: soil biota facilitate knotweed invasion. Ecosphere, 4(2):1–11, 2013.

Parker, I. M. and Gilbert, G. S. The evolutionary ecology of novel plant-pathogen interactions. Annu. Rev. Ecol. Evol. Syst., 35:675–700, 2004.

Pettay, D. T., Wham, D. C., Smith, R. T., Iglesias-Prieto, R., and LaJeunesse, T. C. Microbial invasion of the caribbean by an indo-pacific coral zooxanthella. Proceedings of the National Academy of Sciences, 112(24):7513–7518, 2015.

Piou, D., Delatour, C., and Marçais, B. Hosts and distribution of collybia fusipes in france and factors related to the disease’s severity. Forest Pathology, 32(1):29–41, 2002.

Preston, D. L., Mischler, J. A., Townsend, A. R., and Johnson, P. T. Disease ecology meets ecosystem science. Ecosystems, 19:737–748, 2016.

Prevéy, J. S. and Seastedt, T. R. Increased winter precipitation benefits the native plant pathogen ustilago bullata that infects an invasive grass. Biological Invasions, 17:3041–3047, 2015.

Rohr, R. P., Saavedra, S., and Bascompte, J. On the structural stability of mutualistic systems. Science, 345(6195):1253497, 2014.

Runghen, R., Poulin, R., Monlléo-Borrull, C., and Llopis-Belenguer, C. Network analysis: ten years shining light on host–parasite interactions. Trends in Parasitology, 37(5):445– 455, 2021.

Santini, A., Ghelardini, L., De Pace, C., Desprez-Loustau, M.-L., Capretti, P., Chandelier, A., Cech, T., Chira, D., Diamandis, S., Gaitniekis, T., et al. Biogeographical patterns and determinants of invasion by forest pathogens in europe. New Phytologist, 197(1):238–250, 2013.

Schleuning, M., Fründ, J., and García, D. Predicting ecosystem functions from biodiversity and mutualistic networks: an extension of trait-based concepts to plant–animal interactions. Ecography, 38(4):380–392, 2015.

Schuchert, P., Shuttleworth, C. M., McInnes, C. J., Everest, D. J., and Rushton, S. P. Land-scape scale impacts of culling upon a european grey squirrel population: can trapping reduce population size and decrease the threat of squirrelpox virus infection for the native red squirrel? Biological Invasions, 16:2381–2391, 2014.

Shipunov, A., Newcombe, G., Raghavendra, A. K., and Anderson, C. L. Hidden diversity of endophytic fungi in an invasive plant. American Journal of Botany, 95(9):1096–1108, 2008.

Smith, G. R., Steidinger, B. S., Bruns, T. D., and Peay, K. G. Competition–colonization tradeo?s structure fungal diversity. The ISME journal, 12(7):1758–1767, 2018.

Smith, S. E. and Read, D. J. Mycorrhizal symbiosis. Academic press, 2010.

Strauss, A., White, A., and Boots, M. Invading with biological weapons: the importance of disease-mediated invasions. Functional Ecology, pages 1249–1261, 2012.

Tedersoo, L., Suvi, T., Beaver, K., and Kõoljalg, U. Ectomycorrhizal fungi of the seychelles: diversity patterns and host shifts from the native vateriopsis seychellarum (dipterocarpaceae) and intsia bijuga (caesalpiniaceae) to the introduced eucalyptus robusta (myrtaceae), but not pinus caribea (pinaceae). New phytologist, 175(2):321–333, 2007.

Thrall, P. H., Burdon, J., and Bever, J. D. Local adaptation in the linum marginale—melampsora lini host-pathogen interaction. Evolution, 56(7):1340–1351, 2002.

Vacher, C., Daudin, J.-J., Piou, D., and Desprez-Loustau, M.-L. Ecological integration of alien species into a tree-parasitic fungus network. Biological Invasions, 12:3249–3259, 2010.

Valdovinos, F. S. Mutualistic networks: moving closer to a predictive theory. Ecology Letters, 22(9):1517–1534, 2019.

Van der Putten, W. H. et al. Impacts of soil microbial communities on exotic plant invasions. Trends in Ecology & Evolution, 25(9):512–519, 2010.

Vogelsang, K. M. and Bever, J. D. Mycorrhizal densities decline in association with nonnative plants and contribute to plant invasion. Ecology, 90(2):399–407, 2009.

White, L. A., Forester, J. D., and Craft, M. E. Dynamic, spatial models of parasite transmission in wildlife: Their structure, applications and remaining challenges. Journal of Animal Ecology, 87(3):559–580, 2018.

Wolfe, B. E. and Pringle, A. Geographically structured host specificity is caused by the range expansions and host shifts of a symbiotic fungus. The ISME journal, 6(4):745–755, 2012.

Wu, L., Wang, X.-W., Tao, Z., Wang, T., Zuo, W., Zeng, Y., Liu, Y.-Y., and Dai, L. Data-driven prediction of colonization outcomes for complex microbial communities. Nature Communications, 15(1):2406, 2024.

Yamauchi, A., Nishida, T., and Ohgushi, T. Mathematical model of colonization process of mycorrhizal plants: e?ect of interaction between plants with fungi. Journal of Plant Interactions, 6(2-3):129–132, 2011.

Zenni, R. D. and Nuñez, M. A. The elephant in the room: the role of failed invasions in understanding invasion biology. Oikos, 122(6):801–815, 2013.

