## Supplementary information for "Toward a unified theory of microbially-mediated invasion"

### A Numerical simulations

#### A.1 Overview of interaction scenarios and default parameter values

A brief description of model parameters and their default values used for the simulations is provided in Table A.1. The value of the resource exchange parameters $\alpha$ and $\beta$ for each of the 7 scenarios described in Fig. 2 are provided in Table A.2. Specific parameters used for the plots in Fig. 4 are provided in Table A.3.

| Symbol | Description | Default value |
|---|---|---|
| $p_n$ | Biomass of native host population | – |
| $m_n$ | Biomass of native microbial community | – |
| $p_i$ | Biomass of invasive host population | – |
| $m_i$ | Biomass of invasive microbial community | – |
| $\alpha_{jw}$ | Rate of microbes to hosts resource supply ($j$ to $w$) | 0.4 (0, 0.3) |
| $\beta_{jw}$ | Rate of hosts to microbes resource supply ($j$ to $w$) | 0.4 (0, 0.3) |
| $q_{hp_j}$ | Conversion factor: resources received from microbes into host biomass | 5 |
| $q_{cp_j}$ | Conversion factor: resources supplied to microbes into host biomass | 1 |
| $q_{cm_j}$ | Conversion factor: resources received from hosts into microbial biomass | 1 |
| $q_{hm_j}$ | Conversion factor: resources supplied to hosts into microbial biomass | 1 |
| $\mu_{p_j}$ | Maintenance rate (hosts) | 0.1 |
| $\mu_{m_j}$ | Maintenance rate (microbes) | 0.1 |
| $r_{p_j}$ | Intrinsic growth rate (hosts) | 0.02 |
| $c_{p_{jw}}$ | Competitive effect of host population $j$ on host population $w$ | 0.02 (weak) or 0.12 (strong) |
| $c_{m_{jw}}$ | Competitive effect of microbial community $j$ on microbial community $w$ | 0.02 (weak) or 0.12 (strong) |
| $d$ | Default ratio of host to microbial biomass | 2 |

**Table A.1:** Brief description of model's variables and parameters and their default values used for the simulations. Index $j = n, i$ and $w = n, i$ refer to native ($n$) or invasive ($i$). Values in bracket corresponds to other parameter combinations chosen for the implementation of the scenarios of Fig. 4, as provided in Table A.3. Representative parameters for the interaction scenarios presented in Fig. 2 are provided in Table A.2.

| Interaction Scenario | Parameter values |
|---|---|
| 1. | $\alpha_{in} = 0,\ \beta_{ni} = 0 \quad ;\ \alpha_{ni} = 0,\ \beta_{in} = 0$ |
| 2. | $\alpha_{in} = 0,\ \beta_{ni} = 0 \quad ;\ \alpha_{ni} = \alpha,\ \beta_{in} = \beta$ |
| 3. | $\alpha_{in} = \alpha,\ \beta_{ni} = \beta \quad ;\ \alpha_{ni} = 0,\ \beta_{in} = 0$ |
| 4. | $\alpha_{in} = 0,\ \beta_{ni} = 0 \quad ;\ \alpha_{ni} = 0,\ \beta_{in} = \beta$ |
| 5. | $\alpha_{in} = 0,\ \beta_{ni} = 0 \quad ;\ \alpha_{ni} = \alpha,\ \beta_{in} = 0$ |
| 6. | $\alpha_{in} = 0,\ \beta_{ni} = \beta \quad ;\ \alpha_{ni} = 0,\ \beta_{in} = 0$ |
| 7. | $\alpha_{in} = \alpha,\ \beta_{ni} = 0 \quad ;\ \alpha_{ni} = 0,\ \beta_{in} = 0$ |

**Table A.2:** Possible parameter values of resource exchange rates between native microbes and invasive hosts ($\alpha_{ni}$ and $\beta_{in}$) and between native hosts and invasive microbes ($\alpha_{in}$ and $\beta_{ni}$), for the given interaction scenarios of Fig. 2.

| Figure | Parameter | Value | Figure | Parameter | Value |
|---|---|---|---|---|---|
| 4a | $\alpha_{ii}$ | 0.3 | 4b | $\alpha_{ii}$ | 0.3 |
| | $\beta_{ii}$ | 0.3 | | $\beta_{ii}$ | 0.3 |
| | $\alpha_{ni}$ | 0.4 | | $\alpha_{ni}$ | 0 |
| | $\beta_{in}$ | 0.3 | | $\beta_{in}$ | 0 |
| | $\alpha_{in}$ | 0 | | $\alpha_{in}$ | 0.3 |
| | $\beta_{ni}$ | 0 | | $\beta_{ni}$ | 0.4 |
| | $c_{m_{in}}$ | 0.12 | | $c_{m_{in}}$ | 0.12 |
| | $c_{m_{ni}}$ | 0.02 | | $c_{m_{ni}}$ | 0.02 |
| | $c_{p_{in}}$ | 0.02 | | $c_{p_{in}}$ | 0.12 |
| | $c_{p_{ni}}$ | 0.02 | | $c_{p_{ni}}$ | 0.12 |

**Table A.3:** Brief description of model parameters used to produce Fig. 4. Other parameter values corresponds to those listed in Table A.1.

#### A.2 Scenarios on ModelRxiv

Scenarios of interest are uploaded on the modelRxiv platform, at the following link: `https://modelrxiv.org/model/YfndNX`. The website allows the reproduction and re-parametrization of the 7 interaction scenarios of Fig. 2, as shown in Fig. A.1, and the interactive reproduction of Fig. 4.

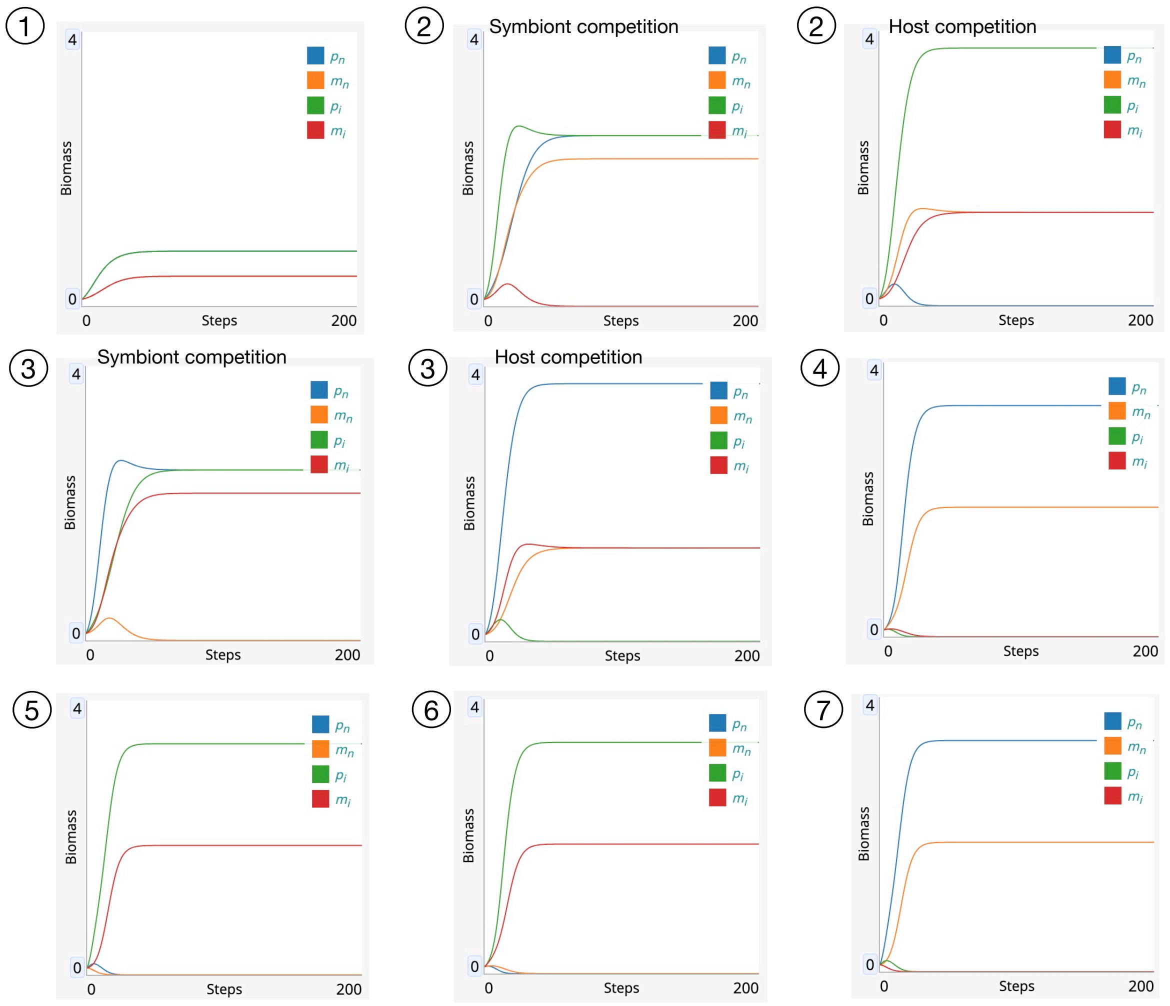

**Fig. A.1:** Timeseries produced by scenarios 1-7 described in Fig. 2, for the parameter combinations provided in Table A.2 and Table A.2. Note that in scenarios 2 and 3 coexistence of hosts or symbionts is unstable when competition is strong, and differences in model parameters or in initial conditions will lead to competitive exclusion of one of the two hosts and one of the two symbionts. The same steady states are stable only for weak competition, as discussed in SI E.1.4.

### B Analysis of the one host and one symbiont case

We here consider the association of one host, with biomass $p$ and intrinsic growth rate $r_p$, that transfers resources to a symbiont at rate $\beta$. The symbiont, with biomass $m$, transfers resources to the host at a rate $\alpha$. The dynamics of their interaction is described by the model of Eq. (3):

$$\begin{cases} \dfrac{dp}{dt} & = r_p p + \dfrac{pm}{\dfrac{p}{d}+m} Q_p(\alpha,\beta) - \mu_p p^2 \,, \\ \dfrac{dm}{dt} & = \dfrac{pm}{\dfrac{p}{d}+m} Q_m(\alpha,\beta) - \mu_m m^2 \,, \end{cases} \quad \text{where} \quad \begin{cases} Q_p(\alpha,\beta) & = q_{hp}\dfrac{\alpha}{d} - q_{cp}\beta \,, \\ Q_m(\alpha,\beta) & = q_{cm}\beta - q_{hm}\dfrac{\alpha}{d} \,. \end{cases} \tag{6}$$

The quantities $Q_p$ and $Q_m$ represent respectively the net gain of the plant and the net gain of the symbiont.

#### B.1 Steady state existence

The system of Eq. (6) presents three steady states: The extinction steady state $(0,0)$, the symbiont-free steady state $(p_0,0)$ with $p_0 = r_p/\mu_p > 0$, and the coexistence steady state $(p^*,m^*)$, which is observed as long as $Q_p + r_p > 0$ and $Q_m > 0$. The positive coexistence steady state $(p^*,m^*)$ is given by

$$p^* = p_0 + \frac{\dfrac{Q_p Q_m}{\mu_m \mu_p}}{\dfrac{S}{d} + \dfrac{Q_m}{\mu_m}} \,, \text{ and } m^* = \frac{d\,Q_m}{\mu_m} \frac{\dfrac{S}{d}}{\dfrac{S}{d} + \dfrac{Q_m}{\mu_m}} \tag{7}$$

where $S$ is the total biomass, $S = p^*/d + m^*$ and it satisfies

$$S = \frac{p_0}{2d} + \sqrt{\left(\frac{p_0}{2d}\right)^2 + p_0\frac{Q_m}{\mu_m} + \frac{Q_m\,Q_p}{\mu_m\,\mu_p}} \,.$$

The phase plane of the system of Eq. 6, with corresponding steady states and nullclines is shown in Fig. B.1.

#### B.2 Parasitic/mutualistic hosts and symbionts

The description of the parasitic/mutualistic interaction are summarized in Fig. 2 and explained in detail in Fig. B.2.

**Mutualistic/parasitic symbiont.** When $Q_p > 0$, that is for

$$\alpha \geq q_{cp}/q_{hp}\beta\, d,$$

the growth rate of the host in association with the symbiont is always larger than the biomass of the host alone ($p_0$). In this case the symbiont is *mutualistic* to the host.

Conversely, when $Q_p < 0$, that is $0 < \alpha \leq q_{cp}/q_{hp}\beta\, d$, the growth rate of the host is reduced compared to its growth rate alone. In this case we say that the symbiont is *parasitic*, and If the intrinsic growth rate of the host in the absence of the symbiont is low, the parasitic symbiont can drive the host to extinction. This occurs for

$$\left(\frac{p_0}{2d}\right)^2 + p_0\frac{Q_m}{\mu_m} + \frac{Q_m\,Q_p}{\mu_m\,\mu_p} < 0.$$

**Mutualistic/parasitic host.** Similarly, if $Q_m > 0$, that is $\beta$ is large enough, such that

$$\beta \geq q_{hm}/q_{cm}\alpha/d,$$

the biomass of the symbiont is positive and the host is *mutualistic* to the symbiont. While if $Q_m \leq 0$, that is $\beta \leq q_{hm}/q_{cm}\alpha/d$, the positive steady state does not exists and the host is *parasitic* to the symbiont. In this case, the host drives the symbiont toward extinction, as symbionts in this model are obligate mutualists.

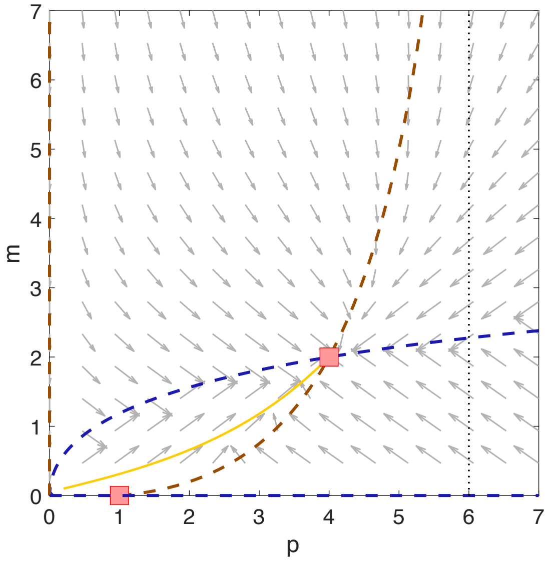

**Fig. B.1:** (a) Phase plane corresponding to the system of equations (6), for which $r_p > 0$. Brown dashed lines are nullclines found for $dp/dt = 0$, while blue dashed lines are nullclines found for $dm/dt = 0$. Intersection of the two non-zero nullclines corresponds to the stable steady state $(p^*, m^*)$, represented by the red square. Intersection of the $dp/dt = 0$ nullcline with the horizontal axis corresponds to the steady state $(p_0, 0)$, in which the host reaches a symbiont-free steady state. The vertical black dotted line corresponds to the asymptote $p = Q_p/\mu_p$.

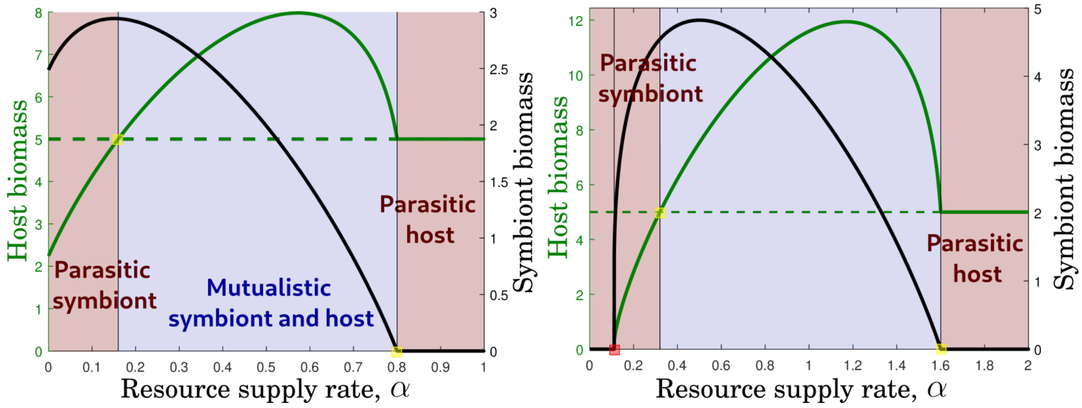

**Fig. B.2:** Host (green plain curve) and symbiont (black plain curve) biomass at equilibrium $(p^*, m^*)$ defined by (7), with respect to the resource exchange rate $\alpha$ of the symbiont. The host exchange rate $\beta$ is fixed to $\beta = 0.4$ (left panel) and $\beta = 0.8$ (right panel). The dashed line corresponds to the biomass of the host alone $p_0 = r_p/\mu_p$, with $r_p = 0.5$. The yellow squares correspond to the critical values when symbiont becomes parasitic, i.e., when $\alpha = q_{cp}/q_{hp}\beta\, d$ (left square), and when host becomes parasitic (right square), i.e., when $\alpha = \beta d q_{cm}/q_{hm}$ The red square corresponds to the threshold when parasitic symbiont become pathogenic, meaning that the host can not grow in the presence of the symbiont.

#### B.3 Steady state stability

To derive the stability of the steady state $(p^*, m^*)$, we compute the Jacobian of the system of equation (6). We obtain:

$$J(p^*, m^*) = \frac{1}{\frac{p^*}{d} + m^*} \begin{pmatrix} \frac{{m^*}^2}{\frac{p^*}{d} + m^*} Q_p + (r_p - 2\mu_p p^*)\left(\frac{p^*}{d} + m^*\right) & \frac{\frac{{p^*}^2}{d}}{\left(\frac{p^*}{d} + m^*\right)} Q_p \\ \frac{{m^*}^2}{\left(\frac{p^*}{d} + m^*\right)} Q_m & \frac{\frac{{p^*}^2}{d}}{\frac{p^*}{d} + m^*} Q_m - 2\mu_m m^* \left(\frac{p^*}{d} + m^*\right) \end{pmatrix}. \quad (8)$$

From Eq. (6), we know that at steady state

$$\begin{cases} \frac{m^*}{\frac{p^*}{d} + m^*} Q_p = \mu_p p^* - r_p = \mu_p (p^* - p_0) \\ \frac{p^*}{\frac{p^*}{d} + m^*} Q_m = \mu_m m^* . \end{cases}$$

Thus the Jacobian $J$ can be written as

$$J(p^*, m^*) = \frac{1}{\frac{p^*}{d} + m^*} \begin{pmatrix} -\mu_p p^* \left(m^* + \frac{2p^* - p_0}{d}\right) & \mu_m m^* \frac{p^*}{d} \frac{Q_p}{Q_m} \\ \mu_p m^* \frac{Q_m}{Q_p} \left(p^* - p_0\right) & -\mu_m m^* \left(2m^* + \frac{p^*}{d}\right) \end{pmatrix}.$$

First, we have $\mathrm{Tr}(J) < 0$. If $Q_p \geq 0$, then $p^* > p_0$ and the inequality follows. Conversely, if $Q_p < 0$ we have

$$\begin{aligned} \mathrm{Tr}(J) &= -\mu_p p^* \left(m^* + \frac{2p^* - p_0}{d}\right) - \mu_m m^* \left(2m^* + \frac{p^*}{d}\right) \\ &= -(\mu_p p^* + \mu_m m^*) \left(m^* + \frac{p^*}{d}\right) - \mu_p p^* \frac{p^* - p_0}{d} - \mu_m m^{*2} \\ &= -(\mu_p p^* + \mu_m m^*) \left(m^* + \frac{p^*}{d}\right) - \frac{p^*}{d} \frac{m^*}{\frac{p^*}{d} + m^*} Q_p - m^* \frac{p^*}{\frac{p^*}{d} + m^*} Q_m < 0 . \end{aligned}$$

The inequality holds true if $Q_m > -Q_p/d$, that is equivalent to $q_{cm} > q_{cp}/d$ and $q_{hm} < q_{hp}/d$, which imposes that

$$\frac{q_{hp} q_{cm}}{q_{cp} q_{hm}} > 1.$$

To prove stability of $(p^*, m^*)$ we should therefore show that $\mathrm{Det}J > 0$. We have

$$\begin{aligned} \mathrm{Det}J &= \mu_m \mu_p m^* p^* \left(m^* + \frac{2p^* - p_0}{d}\right) \left(2m^* + \frac{p^*}{d}\right) - \mu_p \mu_m m^{*2} \frac{p^*}{d} (p^* - p_0) \\ &= \mu_m \mu_p m^* p^* \left(m^* + \frac{p^*}{d}\right) \left(2\left(m^* + \frac{p^*}{d}\right) - \frac{p_0}{d}\right) > 0 . \end{aligned}$$

The positivity of the determinant follows from the following inequality

$$\begin{aligned} 2\left(m^* + \frac{p^*}{d}\right) - \frac{p_0}{d} &= 2S_0 - \frac{p_0}{d} \\ &= \sqrt{\left(\frac{p_0}{2d}\right)^2 + \frac{Q_m}{\mu_m} p_0 + \frac{Q_m \, Q_p}{\mu_m \mu_p}} > 0 . \end{aligned}$$

Therefore, we showed that when the unique positive stable steady state $(p^*, m^*)$ exists, that is when $r_p \geq 0$, $Q_m > 0$ and $\left(\frac{p_0}{2d}\right)^2 + p_0\frac{Q_m}{\mu_m} + \frac{Q_m\,Q_p}{\mu_m\,\mu_p} \geq 0$, then it is stable. The temporal dynamics of $p$ and $m$ over time is shown in Fig. B.1b.

However, when $Q_m < 0$, then the free-symbiont steady state $(p_0, 0)$ becomes stable because its Jacobian is

$$J(p_0, 0) = \begin{pmatrix} -r_p & d\,Q_p \\ 0 & d\,Q_m \end{pmatrix}.$$

### C Analysis of the one host and two symbionts case

We consider the case in which one native host $p_n$ is associated with 2 symbionts (the native symbiont $m_n$ and the invasive symbiont $m_i$) which compete between each other with competition strength $c$. The corresponding differential equation system, is given by:

$$\begin{cases} \dfrac{dp_n}{dt} &= r_p\,p_n + \dfrac{p_n}{\dfrac{p_n}{d} + m_n + m_i}\left(Q_{p_n}m_n + Q_{p_i}m_i\right) - \mu_p p_n^2\,, \\ \dfrac{dm_n}{dt} &= \dfrac{Q_{m_n}p_n m_n}{\dfrac{p_n}{d} + m_n + m_i} - \mu_m m_n^2 - c m_n\,m_i\,, \\ \dfrac{dm_i}{dt} &= \dfrac{Q_{m_i}p_n m_i}{\dfrac{p_n}{d} + m_n + m_i} - \mu_m m_i^2 - c m_n\,m_i\,. \end{cases}$$

The host is either obligate or facultative mutualists ($r_p \geq 0$). The native host and the native symbionts are mutualistic, while the native host and the invasive symbiont might be either parasitic or mutualistic. In particular their exchange rates are given by

$$\begin{cases} Q_{p_n} = q_{hp}\dfrac{\alpha_{nn}}{d} - q_{cp}\beta_{nn} > 0 \\ Q_{m_n} = q_{cm}\beta_{nn} - q_{hm}\dfrac{\alpha_{nn}}{d} > 0 \end{cases} \quad \text{and} \quad \begin{cases} Q_{p_i} = q_{hp}\dfrac{\alpha_{in}}{d} - q_{cp}\beta_{ni}\,, \\ Q_{m_i} = q_{cm}\beta_{ni} - q_{hm}\dfrac{\alpha_{in}}{d}\,. \end{cases}$$

#### C.1 Steady state existence

To compute the steady state, we define the two following vectors $\mathbf{Q}_p$, $\mathbf{Q}_m$ of net gain and the competition matrix $\mathbf{C}_m$

$$\mathbf{Q}_m = \begin{pmatrix} Q_{m_n} \\ Q_{m_i} \end{pmatrix} \quad \text{and} \quad \mathbf{Q}_p = \begin{pmatrix} Q_{p_n} & Q_{p_i} \end{pmatrix} \quad \text{and} \quad \mathbf{C}_m = \begin{pmatrix} \mu_m & c \\ c & \mu_m \end{pmatrix},$$

and the two quantities: the free-symbiont equilibrium of the host $p_0$ and the total biomass of the system $S$;

$$p_0 = \frac{r_p}{\mu_p} \quad \text{and} \quad S = \frac{p}{d} + m_n + m_i.$$

The competition matrix can be reformulate as follows $\mathbf{C}_m = (\mu_m - c)I + c\mathbf{1}$, where $\mathbf{1}$ is the matrix full of 1 and $I$ is the identity matrix. Thus the matrix $\mathbf{C}_m$ is invertible if and only if

$$\mu_m \neq c.$$

and its inverse satisfies

$$\mathbf{C}_m^{-1} = \frac{1}{\mu_m - c}I - \frac{c}{(\mu_m - c)(\mu_m + c)}\mathbf{1}\,.$$

**Coexistence steady state** A positive steady state $(p, m_n, m_i)$, that we write $(p, \mathbf{m})$ with $\mathbf{m} = (m_n, m_i)$, will satisfy the following problem

$$\frac{1}{S}\mathbf{Q}_p\mathbf{m} = \mu_p p - \mu_p p_0 \quad \text{and} \quad \frac{1}{S}\mathbf{Q}_m\, p = \mathbf{C}_m\mathbf{m}\,.$$

By combining the two systems, we obtain

$$\frac{\mathbf{C}_m^{-1}\mathbf{Q}_m\mathbf{Q}_p}{\mu_p}\mathbf{m} = S^2\mathbf{m} - S\mathbf{C}_m^{-1}\mathbf{Q}_m p_0 \quad \text{and} \quad \frac{\mathbf{Q}_p\mathbf{C}_m^{-1}\mathbf{Q}_m}{\mu_p}p = S^2\left(p - p_0\right)\,.$$

We can reformulate the system as follows:

$$p = \frac{S^2\, p_0}{S^2 - \dfrac{\mathbf{Q}_p\mathbf{C}_m^{-1}\mathbf{Q}_m}{\mu_p}} \quad \text{and} \quad (S^2\, I - \mathbf{A})\mathbf{m} = S\mathbf{q}_0\,,$$

where $\mathbf{A} = \dfrac{\mathbf{C}_m^{-1}\mathbf{Q}_m\mathbf{Q}_p}{\mu_p}$ and $\mathbf{q}_0 = \mathbf{C}_m^{-1}\mathbf{Q}_m\, p_0$.

Observing that $\mathrm{Tr}(\mathbf{A}) = \dfrac{\mathbf{Q}_p\mathbf{C}_m^{-1}\mathbf{Q}_m}{\mu_p}$ and $\mathbf{A}^k = \left(\dfrac{\mathbf{Q}_p\mathbf{C}_m^{-1}\mathbf{Q}_m}{\mu_p}\right)^k \mathbf{A}$ for any $k \geq 2$, we deduce that

$$\begin{aligned}(S^2 I - \mathbf{A})^{-1} \quad & \frac{1}{S^2}\left(I - \frac{\mathbf{A}}{S^2}\right)^{-1} = \frac{1}{S^2}\sum\nolimits_{k\geq 0}\left(\frac{\mathbf{A}}{S^2}\right)^k \\ & \frac{1}{S^2}\left(I + \sum\nolimits_{k\geq 1}\left(\frac{1}{S^2}\frac{\mathbf{Q}_p\mathbf{C}_m^{-1}\mathbf{Q}_m}{\mu_p}\right)^k \mathbf{A}\right) \\ &= \frac{1}{S^2}\left(I + \frac{\mathbf{A}}{S^2 - \dfrac{\mathbf{Q}_p\mathbf{C}_m^{-1}\mathbf{Q}_m}{\mu_p}}\right) = \frac{S^2 I + \left(\mathbf{A} - \dfrac{\mathbf{Q}_p\mathbf{C}_m^{-1}\mathbf{Q}_m}{\mu_p}I\right)}{S^2\left(S^2 - \dfrac{\mathbf{Q}_p\mathbf{C}_m^{-1}\mathbf{Q}_m}{\mu_p}\right)}\,.\end{aligned}$$

We can compute $\mathbf{m}$ as follow

$$\mathbf{m} = (S^2 I - \mathbf{A})^{-1} S\,\mathbf{q}_0 = \frac{S^2\mathbf{q}_0 + \left(\mathbf{A} - \dfrac{\mathbf{Q}_p\mathbf{C}_m^{-1}\mathbf{Q}_m}{\mu_p}I\right)\mathbf{q}_0}{S(S^2 - \dfrac{\mathbf{Q}_p\mathbf{C}_m^{-1}\mathbf{Q}_m}{\mu_p})} = \frac{S\mathbf{C}_m^{-1}\mathbf{Q}_m p_0}{S^2 - \dfrac{\mathbf{Q}_p\mathbf{C}_m^{-1}\mathbf{Q}_m}{\mu_p}}\,.$$

Thus from the definition of $S$, we obtain the following equation

$$\begin{aligned} S &= \frac{p}{d} + \mathbf{e}\,\mathbf{m} \\ &= \frac{1}{d}\frac{S^2\, p_0}{S^2 - \dfrac{\mathbf{Q}_p\mathbf{C}_m^{-1}\mathbf{Q}_m}{\mu_p}} + \frac{S\mathbf{e}\mathbf{C}_m^{-1}\mathbf{Q}_m p_0}{S^2 - \dfrac{\mathbf{Q}_p\mathbf{C}_m^{-1}\mathbf{Q}_m}{\mu_p}}\,,\end{aligned}$$

where $\mathbf{e} = (1, \ldots, 1)$. We show that $S$ is the positive root of the following second order polynomial

$$S^2 - \frac{p_0}{d}S - \left(\mathbf{e}\mathbf{C}_m^{-1}\mathbf{Q}_m p_0 + \frac{\mathbf{Q}_p\mathbf{C}_m^{-1}\mathbf{Q}_m}{\mu_p}\right)\,.$$

Finally, using the property of $\mathbf{C}_m^{-1}$, we can show that

$$\mathbf{e}\mathbf{C}_m^{-1}\mathbf{Q}_m = \frac{\mathbf{e}\mathbf{Q}_m}{\mu_m + c} \quad \text{and} \quad \frac{\mathbf{Q}_p\mathbf{C}_m^{-1}\mathbf{Q}_m}{\mu_p} = \frac{\mathbf{Q}_p\mathbf{Q}_m}{\mu_p(\mu_m - c)} - \frac{c\,(\mathbf{e}\mathbf{Q}_m)(\mathbf{e}\mathbf{Q}_p)}{(\mu_m - c)(\mu_m + c)\mu_p}\,,$$

and we deduce that $S$ is given by

$$S = \frac{p_0}{2d} + \sqrt{\left(\frac{p_0}{2d}\right)^2 + \frac{\mathbf{e}\mathbf{Q}_m}{\mu_m + c} p_0 + \frac{\mathbf{Q}_p \mathbf{C}_m^{-1} \mathbf{Q}_m}{\mu_p}}\,.$$

Finally, we get

$$p = p_0 + \frac{\dfrac{\mathbf{Q}_p \mathbf{C}_m^{-1} \mathbf{Q}_m}{\mu_p}}{\dfrac{S}{d} + \dfrac{\mathbf{e}\mathbf{Q}_m}{\mu_m + c}} \quad \text{and} \quad \mathbf{m} = \frac{S}{\dfrac{S}{d} + \dfrac{\mathbf{e}\mathbf{Q}_m}{\mu_m + c}} \mathbf{C}_m^{-1} \mathbf{Q}_m\,. \tag{9}$$

**Coexistence steady exists only under weak competition.** The coexistence steady state is positive if and only if $S$ exists and the components of $\mathbf{m}$ are positive, that is when the following inequality hold true:

$$\left(\frac{p_0}{2d}\right)^2 + \frac{\mathbf{e}\mathbf{Q}_m}{\mu_m + c} p_0 + \frac{\mathbf{Q}_p \mathbf{C}_m^{-1} \mathbf{Q}_m}{\mu_p} > 0, \quad \mu_m Q_{m_n} - c Q_{m_i} > 0 \ \text{ and } \ \mu_m Q_{m_i} - c Q_{m_n} > 0$$

The two last inequalities imposes that

$$c < \mu_m \ \text{ and } \ \frac{\mu_m}{c} Q_{m_n} > Q_{m_i} > \frac{c}{\mu_m} Q_{m_n} > 0.$$

Thus, the coexistence steady state does exist if the **symbionts compete weakly ($c < \mu_m$)**, the host is not *parasitic* to the invasive symbiont ($Q_{m_i} < \mu_m\, Q_m/c$) and the invader symbiont has no *competing advantage* over the native symbiont ($Q_{m_i} > c\, Q_m/\mu_m$).

#### C.2 Parasitic versus mutualistic behaviour of host and symbiont

We here investigate the different nature of the interactions between two symbionts (a native and an invasive one) and a host, when the host is initially associated with a native symbiont, that can competes with the invasive symbiont.

**Parasitic host.** A host associated with a native mutualistic symbiont can be parasitic to an invasive symbiont, that competes with the native symbiont ($Q_{m_i} < \mu_m\, Q_m/c$). The host can be parasitic to the invasive symbiont even if it would have been mutualistic to it in absence of the native symbiont ($Q_{m_i} > 0$). This detrimental effect of the host associated with a native symbiont is due to competition between the two symbionts (see rightmost yellow square in Fig. C.1(b) that shows the transition between mutualistic and parasitic host). Thus the association with a mutualistic symbiont can enhance the parasitic nature of the host with respect to invasive symbionts (compare dashed curve $Q_{m_i} = 0$ with plain curve $Q_{m_i} = \mu_m\, Q_m/c$ in Fig. C.1(a)).

**Pathogenic invasive symbiont.** A invasive symbiont is *pathogenic* if it drives the host to extinction. This situation occurs when the exchange rates satisfies the following conditions

$$\left(\frac{p_0}{2d}\right)^2 + \frac{Q_{m_i}}{\mu_m} p_0 + \frac{Q_{p_i} Q_{m_i}}{\mu_p} < 0$$

The transition between a pathogenic and a parasitic symbiont is illustrated by the leftmost red square in Fig. C.1.

**Substitution of native symbiont with invasive symbiont.** If the invasive symbiont has a competing advantage over the native symbiont, that is $Q_{m_i} < c\,Q_m/\mu_m$, then the coexistence steady state does not exist and the invasive symbiont replaces the native symbiont. In this case, the invasive symbiont has a *competing advantage over the native symbiont.* The transition between symbiont substitution and coexistence is depicted by the rightmost red square in Fig. C.1.

The invasive symbiont can be either parasitic or mutualistic to the native host depending on their exchange rates. For instance in absence of native species, the previous definition of section B.2 states that if $Q_{p_i} > 0$ the invasive symbiont is *mutualistic* to the host, while if $Q_{p_i} > 0$ then the symbiont is also *parasitic* to the host. However, this definition only compare the biomass of the host with and without the invasive symbiont. In presence of initial native symbiont, we may also compare with the host biomass when associated with the mutualistic native symbiont. We discuss the parasitic/mutualistic nature of the invasive symbiont with respect to the native host in the following paragraph.

**Mutualistic/parasitic invasive symbiont.** In the presence of a native symbiont, the mutualistic nature of the interaction between the native host and the invasive symbiont may vary when comparing with the scenario without a native symbiont, that competes with the invasive symbiont. In both situations, the invasive symbiont is *mutualistic* to the host, if the biomass of the native host is larger than the biomass of the native host when alone with or without its native symbiont.

When the native and invasive symbionts persist at equilibrium, that is $Q_{m_i} > c\,Q_m/\mu_m$, the invasive symbiont is *mutualistic* when the exchange rates between the native host and the invasive symbiont are such that

$$\frac{\mu_m \mathbf{Q}_p \mathbf{C}_m^{-1} \mathbf{Q}_m}{S^2} \geq \frac{Q_{p_n} Q_{m_n}}{S_n^2} \quad \text{where} \quad S_n = \frac{p_0}{2d} + \sqrt{\left(\frac{p_0}{2d}\right)^2 + \frac{Q_{m_n}}{\mu_m} p_0 + \frac{Q_{p_n} Q_{m_n}}{\mu_p \mu_m}}\,.$$

The transition between a mutualistic and parasitic invasive symbiont is illustrated by the leftmost yellow square in Fig. C.1.

When the invasive symbiont replaces the native symbiont, that is $Q_{m_i} \leq c\,Q_m/\mu_m$, then the invasive symbiont is mutualistic if the biomass of the host is larger than the biomass of the host when associated with its initial native symbiont. This situation occurs when

$$\frac{Q_{p_i} Q_{m_i}}{S_i^2} \geq \frac{Q_{p_n} Q_{m_n}}{S_n^2} \quad \text{where} \quad S_{n/i} = \frac{p_0}{2d} + \sqrt{\left(\frac{p_0}{2d}\right)^2 + \frac{Q_{m_{n/i}}}{\mu_m} p_0 + \frac{Q_{p_{n/i}} Q_{m_{n/i}}}{\mu_p \mu_m}}\,.$$

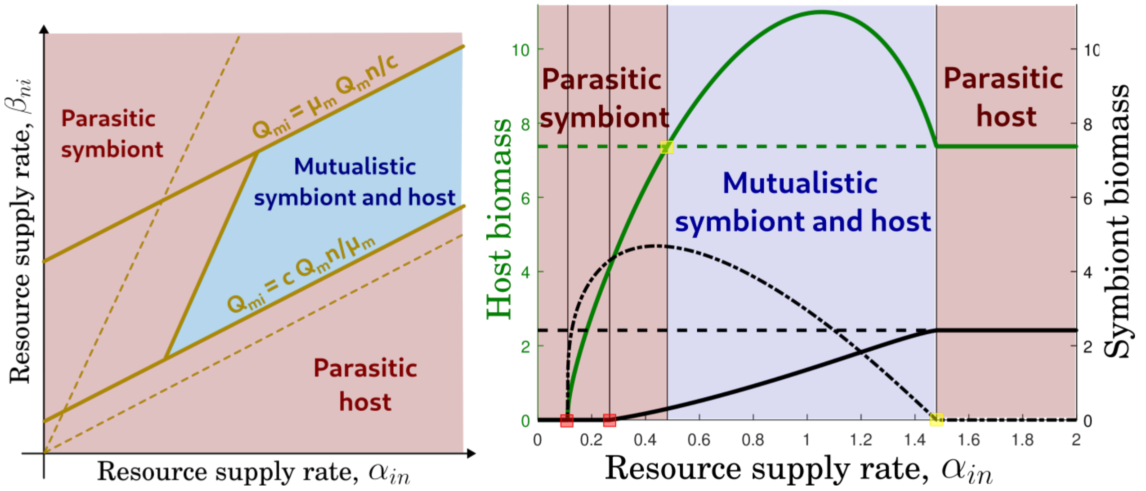

(a) (b)

**Fig. C.1:** Parasitic or mutualistic invasive symbiont and native host. Evolution of the host (green plain curve) and symbiont (black plain curve) biomass at equilibrium defined by (9), with respect to the exchange rate $\alpha_{in}$ of the invasive symbiont to the native host. The exchange rates between native pair are fixed to $\beta_{nn} = \alpha_{nn} = 0.4$. The host exchange rate $\beta_{ni}$ with the invasive symbiont is fixed to $\beta_{ni} = 0.8$. The intrinsic growth rate of the host is fixed to $r_p = 0.5$. The dashed line corresponds to the biomass of the native pair host-symbiont alone. The red square corresponds to the critical values such that either the invader drives the system toward extinction (left square) or the invasive symbiont replaces the native symbiont (right square). The yellow square corresponds to the parasitic mutualistic nature of interactions between native host and invasive symbiont: parasitic symbiont (left square), parasitic host (right square).

### 864 D Analysis of the one symbiont and two competing hosts

We consider the case in which one native symbiont $m$ is associated with 2 hosts (the native host $p_n$ and the invasive host $p_i$) which compete between each other at a rate $c$. The corresponding differential equation system, is given by:

$$\begin{cases} \dfrac{dp_n}{dt} &= r_{p_n} p_n + \dfrac{Q_{p_n} m\, p_n}{\dfrac{p_n}{d} + \dfrac{p_i}{d} + m} - \mu_p p_n^2 - c_{p_n} p_i \,, \\ \dfrac{dp_i}{dt} &= r_{p_i} p_i + \dfrac{Q_{p_i} m\, p_i}{\dfrac{p_n}{d} + \dfrac{p_i}{d} + m} - \mu_p p_i^2 - c_{p_i} p_n \\ \dfrac{dm}{dt} &= \dfrac{m}{\dfrac{p_n}{d} + \dfrac{p_i}{d} + m} \left(Q_{m_n} p_n + Q_{m_i} p_i\right) - \mu_m m^2 \,. \end{cases}$$

The hosts are either obligate or facultative mutualists ($r_{p_n} \geq 0$ and $r_{p_i} \geq 0$). The native host and the native symbionts are mutualistic, while the invasive host and the native symbiont might be either parasitic or mutualistic. In particular their exchange rates are given by

$$\begin{cases} Q_{p_n} = q_{hp}\dfrac{\alpha_{nn}}{d} - q_{cp}\beta_{nn} > 0 \\ Q_{m_n} = q_{cm}\beta_{nn} - q_{hm}\dfrac{\alpha_{nn}}{d} > 0 \end{cases} \quad \text{and} \quad \begin{cases} Q_{p_i} = q_{hp}\dfrac{\alpha_{ni}}{d} - q_{cp}\beta_{in} \,, \\ Q_{m_i} = q_{cm}\beta_{in} - q_{hm}\dfrac{\alpha_{ni}}{d} \,. \end{cases}$$

#### D.1 Steady state existence

To compute the steady state, we define the two following vectors $\mathbf{Q}_p$, $\mathbf{Q}_m$ of net gain and the competition matrix $\mathbf{C}_m$

$$\mathbf{Q}_p = \begin{pmatrix} Q_{p_n} \\ Q_{p_i} \end{pmatrix} \quad \text{and} \quad \mathbf{Q}_m = \begin{pmatrix} Q_{m_n} & Q_{m_i} \end{pmatrix} \quad \text{and} \quad \mathbf{C}_p = \begin{pmatrix} \mu_p & c \\ c & \mu_p \end{pmatrix},$$

The competition matrix can be reformulate as follows $\mathbf{C}_p = (\mu_p - c)I + c\mathbf{1}$, where $\mathbf{1}$ is the matrix full of 1 and $I$ is the identity matrix. Thus the matrix $\mathbf{C}_p$ is invertible if and only if

$$\mu_p \neq c.$$

and its inverse satisfies

$$\mathbf{C}_p^{-1} = \frac{1}{\mu_p - c} I - \frac{c}{(\mu_p - c)(\mu_p + c)} \mathbf{1}\,.$$

We also define the two quantities: the free-symbiont equilibrium of coexisting hosts $\mathbf{p}_0$ and the total biomass of the system $S$;

$$\mathbf{p}_0 = \mathbf{C}_p^{-1} \begin{pmatrix} r_{p_n} \\ r_{p_i} \end{pmatrix} \quad \text{and} \quad S = \frac{p_n}{d} + \frac{p_i}{d} + m.$$

Let us remark that the components of $\mathbf{p}_0$ are all positive, if and only if

$$c_p \leq \mu_p.$$

We immediately recover that if competition between hosts is too strong, coexistence of hosts is not possible.

**Coexistence steady state** A positive steady state $(p_n, p_i, m)$ will satisfy the following problem:

$$\frac{1}{S}\mathbf{Q}_p m = \mathbf{C}_p \mathbf{p} - \mathbf{C}_p \mathbf{p}_0 \quad \text{and} \quad \frac{1}{S}\mathbf{Q}_m \mathbf{p} = \mu_m m\,.$$

Combining the two systems, we obtain:

$$\frac{\mathbf{Q}_m \mathbf{C}_p^{-1} \mathbf{Q}_p}{\mu_m} m = S^2 m - S\frac{\mathbf{Q}_m}{\mu_m}\mathbf{p}_0 \quad \text{and} \quad \frac{\mathbf{C}_p^{-1}\mathbf{Q}_p\mathbf{Q}_m}{\mu_m}\mathbf{p} = S^2\left(\mathbf{p} - \mathbf{p}_0\right).$$

We can reformulate the system as follows:

$$(S^2 I - \mathbf{A})\mathbf{p} = S^2 \mathbf{p}_0 \quad \text{and} \quad m = \frac{S q_0}{S^2 - \dfrac{\mathbf{Q}_m \mathbf{C}_p^{-1} \mathbf{Q}_p}{\mu_m}},$$

where $\mathbf{A} = \dfrac{\mathbf{C}_p^{-1}\mathbf{Q}_p\mathbf{Q}_m}{\mu_m}$, and $q_0 = \dfrac{\mathbf{Q}_m}{\mu_m}\mathbf{p}_0$.

Observing that $\mathrm{Tr}(\mathbf{A}) = \dfrac{\mathbf{Q}_m \mathbf{C}_p^{-1} \mathbf{Q}_p}{\mu_m}$ and $\mathbf{A}^k = \left(\dfrac{\mathbf{Q}_m \mathbf{C}_p^{-1} \mathbf{Q}_p}{\mu_m}\right)^{k-1} \mathbf{A}$ for any $k \geq 2$, we deduce that

$$\begin{aligned}(S^2I-\mathbf{A})^{-1} &= \frac{1}{S^2}\left(I-\frac{\mathbf{A}}{S^2}\right)^{-1} = \frac{1}{S^2}\sum_{k\geq 0}\left(\frac{\mathbf{A}}{S^2}\right)^k \\ &= \frac{1}{S^2}\left(I+\sum_{k\geq 1}\left(\frac{1}{S^2}\frac{\mathbf{Q}_m\mathbf{C}_p^{-1}\mathbf{Q}_p}{\mu_m}\right)^k\mathbf{A}\right) \\ &= \frac{1}{S^2}\left(I+\frac{\mathbf{A}}{S^2-\dfrac{\mathbf{Q}_m\mathbf{C}_p^{-1}\mathbf{Q}_p}{\mu_m}}\right) = \frac{S^2I+\left(\mathbf{A}-\dfrac{\mathbf{Q}_m\mathbf{C}_p^{-1}\mathbf{Q}_p}{\mu_m}I\right)}{S^2\left(S^2-\dfrac{\mathbf{Q}_m\mathbf{C}_p^{-1}\mathbf{Q}_p}{\mu_m}\right)}.\end{aligned}$$

We can compute $\mathbf{p}$ as follows

$$\mathbf{p} = (S^2I-\mathbf{A})^{-1}S^2\mathbf{p}_0 = \frac{S^2\mathbf{p}_0+\left(\mathbf{A}-\dfrac{\mathbf{Q}_m\mathbf{C}_p^{-1}\mathbf{Q}_p}{\mu_m}I\right)\mathbf{p}_0}{\left(S^2-\dfrac{\mathbf{Q}_m\mathbf{C}_p^{-1}\mathbf{Q}_p}{\mu_m}\right)}.$$

Using the definition of $S$, we show that $S$ is the positive root of the following third order polynomial

$$S^3-\frac{\mathbf{e}\mathbf{p}_0}{d}S^2-\left(\frac{\mathbf{Q}_m}{\mu_m}\mathbf{p}_0+\frac{\mathbf{Q}_m\mathbf{C}_p^{-1}\mathbf{Q}_p}{\mu_m}\right)S-\mathbf{e}\left(\frac{\mathbf{C}_p^{-1}\mathbf{Q}_p\mathbf{Q}_m}{\mu_m}-\frac{\mathbf{Q}_m\mathbf{C}_p^{-1}\mathbf{Q}_p}{\mu_m}I\right)\frac{\mathbf{p}_0}{d}.$$

#### D.2 Parasitic versus mutualistic behaviour of host and symbiont

We here investigate the different nature of the interactions between two hosts (a native and an invasive one) and a symbiont, when the symbiont is initially associated with a native host, that can competes with the invasive host.

**Parasitic symbiont.** A symbiont associated with a native mutualistic host can be parasitic or pathogenic to an invasive host, that competes with the native host. It occurs when the exchange rate $\beta_{in}$ is too large (rightmost yellow square Fig. D.2).

**Pathogenic host** If the host is too parasitic, $Q_{m_i}+Q_{m_n}<0$ that corresponds to a low $\beta_{in}$, then invasive host drives the native symbiont toward extinction (leftmost red square in Fig.D.2).

**Mutualistic/parasitic invasive host.** When exchange rate $\beta_{in}$ increases but remains low, then the invasive host is still parasitic with the symbiont as its biomass remains low until it becomes larger than its biomass at equilibrium with its native host alone. The mutualistic/parasitic threshold is represented by the leftmost yellow square in Fig; D.2.

**Substitution of native host with invasive host.** The invasive host may replace the native host when it gain competitive advantage from the native symbiont. It corresponds to the region between the two rightmost red square in Fig.D.2.

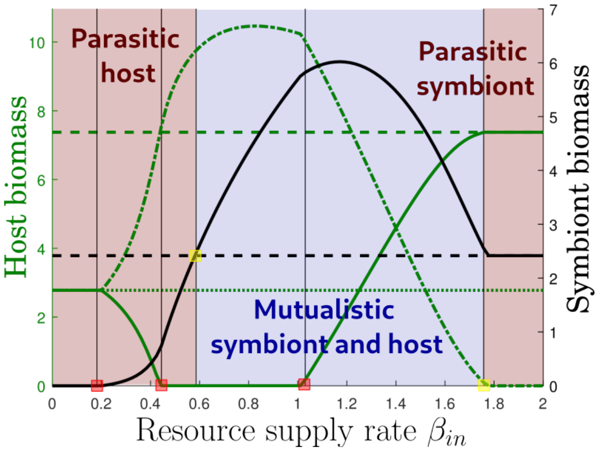

**Fig. D.2:** Parasitic or mutualistic invasive host and native symbiont. Evolution of the native host (solid green curve), invasive host (dashed green curve) and symbiont (solid black curve) biomass at equilibrium defined by (9), with respect to the exchange rate $\beta_{in}$ of the invasive host to the native symbiont. The exchange rates between native pair are fixed to $\beta_{nn} = \alpha_{nn} = 0.4$. The symbiont exchange rate $\alpha_{ni}$ with the invasive host is fixed to $\alpha_{ni} = 0.8$. The host intrinsic growth rates are fixed to $r_p = 0.5$. The dashed line corresponds to the biomass of the native host-symbiont pair alone. The red squares correspond to the critical values such that the invader host drives the symbiont toward extinction (leftmost square), the invasive host replaces the native host (between the middle and rightmost squares). The yellow squares correspond to the parasitic mutualistic nature of interactions between native host and invasive symbiont: parasitic host (left square), parasitic symbiont (right square).

### E Analysis of the different interaction scenarios

We consider here the main system (1) consisting of a native host associated with its native symbiont and an invasive host with its invasive symbiont. As in the main text, we assume that both the native host and the invasive host exchange nutrients with their respective symbionts at similar rates, i.e., $\alpha_{nn} = \alpha_{ii} = \alpha > 0$ and $\beta_{nn} = \beta_{ii} = \beta > 0$. Furthermore, we assume that the native pair and the invasive pair are mutualistic, that is their net gain $Q_m$ and $Q_p$ are positive:

$$Q_p = q_{hp}\frac{\alpha}{d} - q_{cp}\beta > 0 \quad \text{and} \quad Q_m = q_{cm}\beta - q_{hm}\frac{\alpha}{d} > 0\,.$$

In contrast, the exchange between native and invasive species may vary depending on the scenario: (1) no interactions between invasive and native species, (2)-(3) mutualistic interactions between invasive and native species, (4)-(7) parasitic interactions between invasive and native species (either parasitic host or parasitic symbiont), (see Table A.2 and Fig. 3 for details on the scenarios). The net gain resulting from the interaction between the native and invasive $Q_m$ and $Q_p$ will take the following values depending on the scenario

$$Q_p = \begin{cases} 0 & \text{neutral } (1) \\ q_p^- = -q_{cp}\beta < 0 & \text{parasitic symbiont } (4)-(6) \\ Q_p = q_{hp}\dfrac{\alpha}{d} - q_{cp}\beta > 0 & \text{mutualistic host and symbiont } (2)-(3) \\ q_p^+ = q_{hp}\dfrac{\alpha}{d} > Q_p > 0 & \text{parasitic host } (5)-(7) \end{cases}$$

and

$$Q_m = \begin{cases} 0 & \text{neutral } \textcircled{1} \\ q_m^+ = q_{cm}\beta > Q_m > 0 & \text{parasitic symbiont } \textcircled{4} - \textcircled{6} \\ Q_m = q_{cm}\beta - q_{hm}\dfrac{\alpha}{d} > 0 & \text{mutualistic host and symbiont } \textcircled{2} - \textcircled{3} \\ q_m^- = -q_{hm}\dfrac{\alpha}{d} < 0 & \text{parasitic host } \textcircled{5} - \textcircled{7} \end{cases}$$

The model of Eq. (1) can be written as:

$$\begin{cases} \dfrac{dp_n}{dt} &= r_{p_n} p_n + \dfrac{p_n}{\dfrac{p_n}{d} + \dfrac{p_i}{d} + m_n + m_i} \left(m_n Q_p + m_i Q_{p_{ni}}\right) - \mu_p p_n^2 - c_p p_i p_n \,, \\ \dfrac{dm_n}{dt} &= \dfrac{m_n}{\dfrac{p_n}{d} + \dfrac{p_i}{d} + m_n + m_i} \left(p_n Q_m + p_i Q_{m_{in}}\right) - \mu_m m_n^2 - c_m m_i m_n \,, \\ \dfrac{dp_i}{dt} &= r_{p_i} p_i + \dfrac{p_i}{\dfrac{p_n}{d} + \dfrac{p_i}{d} + m_n + m_i} \left(m_i Q_p + m_n Q_{p_{in}}\right) - c_p p_n p_i - \mu_p p_i^2 \,, \\ \dfrac{dm_i}{dt} &= \dfrac{m_i}{\dfrac{p_n}{d} + \dfrac{p_i}{d} + m_n + m_i} \left(p_i Q_m + p_n Q_{m_{ni}}\right) - c_m m_n m_i - \mu_m m_i^2 \,. \end{cases} \tag{10}$$

For our analysis, it will be useful to define the two exchange matrices $\mathbf{Q}_p$ and $\mathbf{Q}_m$:

$$\mathbf{Q}_p = \begin{pmatrix} Q_p & Q_{p_{ni}} \\ Q_{p_{in}} & Q_p \end{pmatrix} \quad \text{and} \quad \mathbf{Q}_m = \begin{pmatrix} Q_m & Q_{m_{in}} \\ Q_{m_{ni}} & Q_m \end{pmatrix} \,,$$

One can remark that the interactions between the native host and the invasive symbiont are described by the couple $(Q_{p_{ni}}, Q_{m_{ni}})$, while the interaction between the invasive host and the native symbiont corresponds to $(Q_{p_{in}}, Q_{m_{in}})$.

We also define the two competition matrices $\mathbf{C}_p$ and $\mathbf{C}_m$:

$$\mathbf{C}_p = \begin{pmatrix} \mu_p & c_p \\ c_p & \mu_p \end{pmatrix} \quad \text{and} \quad \mathbf{C}_m = \begin{pmatrix} \mu_m & c_m \\ c_m & \mu_m \end{pmatrix} \,,$$

and the quantity $S = \dfrac{p_n}{d} + \dfrac{p_i}{d} + m_n + m_i$.

Then, using the notation $\mathbf{p} = (p_n, p_i)$, $\mathbf{m} = (m_n, m_i)$, and $\mathbf{r}_p = (r_{p_n}, r_{p_i})$, the system (10) can be reformulated as

$$\mathbf{p}' = \mathbf{p} \cdot \left(\mathbf{r}_p + \frac{\mathbf{Q}_p \mathbf{m}}{S} - \mathbf{C}_p \mathbf{p}\right)$$
$$\mathbf{m}' = \mathbf{m} \cdot \left(\frac{\mathbf{Q}_m \mathbf{p}}{S} - \mathbf{C}_m \mathbf{m}\right) \,.$$

#### E.1 Steady states of the system

In order to understand the outcome of the interactions between native and invasive species, we first describe the possible steady states of the system.

##### E.1.1 Exclusion steady states consisting of one host and one symbiont

In our system, the steady state that comprises only one host and one symbiont always exists. Therefore, there are four possible exclusion steady states:

$$\begin{array}{ll} (p_n^*, m_n^*, 0, 0) & \text{native pair} \\ (0, 0, p_i^*, m_i^*) & \text{invasive pair} \\ (p_n^*, 0, 0, m_i^*) & \text{mixed pair, invasive symbiont with native host} \\ (0, m_n^*, p_i^*, 0) & \text{mixed pair, invasive host with native symbiont} \end{array} \tag{11}$$

Due to our choice of parameters (i.e., with $\alpha_{nn}$, $\alpha_{ii}$, $\beta_{nn}$ and $\beta_{ii}$ large enough, such that the association between $p_n$ and $m_n$ and $p_i$ and $m_i$ is mutualistic), the steady state with only native species or only invasive species always exists. However, the mixed steady states composed of either a native host with an invasive symbiont, or an invasive host with a native symbiont, exist only when cross-species interactions between $p_n$ and $m_i$ and between $p_i$ and $m_i$ are mutualistic, as shown in scenarios ②-③ (see section B.1).

###### E.1.2 Inclusion steady states consisting of one host and two symbionts

This case corresponds to the situation in which a novel association between a host and a new symbiont of another host is observed, where the host with two symbionts successively outcompete the host. There are two possible steady states of this form :

$$(p_n, 0, m_n, m_i) \text{ and } (0, p_i, m_n, m_i) \text{ symbiont inclusion state.}$$

These equilibria exist if **competition between the symbionts is weak** and the symbionts are mutualistic to the host (see section C.1).

###### E.1.3 Inclusion steady states consisting of two hosts and one symbiont

This inclusion steady state corresponds to the case where one symbiont associates with two hosts, and excludes the other symbiont. In our system, we have two possible steady states of this form

$$(p_n, p_i, m_n, 0) \text{ and } (p_n, p_i, 0, m_i) \text{ host inclusion state.}$$

These equilibria exist if **competition between hosts is weak** and the hosts are mutualistic to the symbionts (see section D.1).

###### E.1.4 Coexistence steady state consisting of two hosts and two symbionts

The last possible equilibrium is the coexistence state where native and invasive species survive, that i s

$$(p_n, p_i, m_n, m_i) \text{ coexistence state.}$$

This steady state only exists when **competition between symbionts and between hosts is weak**.

#### E.2 Outcome of the different interaction scenarios

We first investigate the outcome of the 7 scenarios under strong competition between symbionts and between hosts, that is

$$c_p > \mu_p \text{ and } c_m > \mu_m. \tag{12}$$

Then, we discuss scenarios ② and ③, that correspond to mutualistic host-symbiont interactions under weak competition to show that inclusion steady state or even coexistence can occur.

###### E.2.1 Strong competition between hosts and symbionts

In this situation, only the exclusion equilibrium, defined in (11) can exist and their stability crucially depends on the interactions. To define this dependence more precisely, we compute the Jacobian matrix of the steady state. For any of the four exclusion equilibrium, the Jacobian takes the following form

$$J = \left( \begin{array}{c|c} J(p^*, m^*) & B \\ \hline 0 & I \end{array} \right),$$

where $p^*$, $m^*$ stands for the positive biomass of host an symbiont at equilibrium, $J(p^*, m^*)$ is defined by (8) in section B.3 and $B$ and $I$ are $2 \times 2$ matrices, and $I$ describes the interactions between the excluded host and symbiont and the surviving ones. From the analysis of Section B.3, we know that the matrix $J(p^*, m^*)$ has two negative eigenvalues. The matrix $I$ is diagonal with coefficients that depend on the exclusion equilibrium as follows

**Native pair** $(p_n, m_n, 0, 0)$

$$I_{nn} = \begin{pmatrix} \dfrac{p_0}{S^2 - \dfrac{Q_p Q_m}{\mu_p \mu_m}} \left( \left(1 - \dfrac{c_p}{\mu_p}\right) S^2 - \dfrac{Q_p Q_m}{\mu_p \mu_m} \left(1 - \dfrac{Q_{p_{in}}}{Q_p}\right) \right) & 0 \\ 0 & \left( \dfrac{Q_{m_{ni}}}{Q_m} \mu_m - c_m \right) m_n . \end{pmatrix}$$

**Invasive pair** $(0, 0, p_i, m_i)$

$$I_{ii} = \begin{pmatrix} \dfrac{p_0}{S^2 - \dfrac{Q_p Q_m}{\mu_p \mu_m}} \left( \left(1 - \dfrac{c_p}{\mu_p}\right) S^2 - \dfrac{Q_p Q_m}{\mu_p \mu_m} \left(1 - \dfrac{Q_{p_{ni}}}{Q_p}\right) \right) & 0 \\ 0 & \left( \dfrac{Q_{m_{in}}}{Q_m} \mu_m - c_m \right) m_i \end{pmatrix}.$$

**Mixed pair, invasive symbiont spillover,** $(p_n, 0, m_i, 0)$

$$I_{ni} = \begin{pmatrix} \dfrac{p_0}{S^2 - \dfrac{Q_{p_{ni}} Q_{m_{ni}}}{\mu_p \mu_m}} \left( \left(1 - \dfrac{c_p}{\mu_p}\right) S^2 - \dfrac{Q_{p_{ni}} Q_{m_{ni}}}{\mu_p \mu_m} \left(1 - \dfrac{Q_p}{Q_{p_{ni}}}\right) \right) & 0 \\ 0 & \left( \dfrac{Q_m}{Q_{m_{ni}}} \mu_m - c_m \right) m_i \end{pmatrix}.$$

**Mixed pair, invasive host invasion,** $(0, m_n, p_i, 0)$

$$I_{in} = \begin{pmatrix} \dfrac{p_0}{S^2 - \dfrac{Q_{p_{in}} Q_{m_{in}}}{\mu_p \mu_m}} \left( \left(1 - \dfrac{c_p}{\mu_p}\right) S^2 - \dfrac{Q_{p_{in}} Q_{m_{in}}}{\mu_p \mu_m} \left(1 - \dfrac{Q_p}{Q_{p_{in}}}\right) \right) & 0 \\ 0 & \left( \dfrac{Q_m}{Q_{m_{ni}}} \mu_m - c_m \right) m_n \end{pmatrix}$$

(1) **Native and invasive species do not share symbionts nor hosts.** In this scenario, the cross-species exchange are null, that is $Q_{p_{in}} = Q_{p_{ni}} = 0 = Q_{m_{in}} = Q_{m_{ni}}$, and the exchange matrices $\mathbf{Q}_m$ and $\mathbf{Q}_p$ become

$$\mathbf{Q}_p = \begin{pmatrix} Q_p & 0 \\ 0 & Q_p \end{pmatrix} \quad \text{and} \quad \mathbf{Q}_m = \begin{pmatrix} Q_m & 0 \\ 0 & Q_m \end{pmatrix},$$

Under strong competition (see condition Eq. (12)), we deduce that the matrices $I_{nn}$ and $I_{ii}$ have negative coefficients, and the **exclusion steady states with either native or invasive species are stable**.

**② The association between native symbionts and invasive hosts is mutualistic.** In this scenario, association between native symbionts and invasive hosts is mutualistic, that is $Q_{p_{ni}} = Q_p > 0$ and $Q_{m_{ni}} = Q_m > 0$, while there is no association between native hosts and invasive symbionts, that is $Q_{p_{in}} = 0 = Q_{m_{in}}$. The exchange matrices $\mathbf{Q}_m$ and $\mathbf{Q}_p$ become

$$\mathbf{Q}_p = \begin{pmatrix} Q_p & 0 \\ Q_p & Q_p \end{pmatrix} \text{ and } \mathbf{Q}_m = \begin{pmatrix} Q_m & Q_m \\ 0 & Q_m \end{pmatrix},$$

Under strong competition, we deduce that the matrices $I_{nn}$, $I_{ii}$ and $I_{ni}$ have negative coefficients, and the **exclusion steady states with either native or invasive species and the mixed steady state of invasive hosts and native symbionts are stable**.

**③ The association native hosts and invasive symbionts is mutualistic.** Similarly to previous scenario, the exchange between invasive symbiont and native host are positive, that is $Q_{p_{in}} = Q_p > 0$ and $Q_{m_{in}} = Q_m > 0$, while the exchange between invasive host and native symbiont are null, that is $Q_{p_{ni}} = 0 = Q_{m_{ni}}$. The exchange matrices $\mathbf{Q}_m$ and $\mathbf{Q}_p$ become

$$\mathbf{Q}_p = \begin{pmatrix} Q_p & Q_p \\ 0 & Q_p \end{pmatrix} \text{ and } \mathbf{Q}_m = \begin{pmatrix} Q_m & 0 \\ Q_m & Q_m \end{pmatrix},$$

Under strong competition (Eq. (12)), we deduce that matrices $I_{nn}$, $I_{ii}$ and $I_{in}$ have negative coefficients, and the **exclusion steady states with either native or invasive species and the mixed pair with native host and invasive symbiont are stable**.

**④ Native symbionts are parasitic to invasive hosts.** In this scenario, native symbionts are parasitic to invasive hosts, that is $Q_{p_{in}} = q_p^- < 0$ and $Q_{m_{in}} = q_m^+ > Q_m > 0$, while there is no association between native host and invasive symbiont, that is $Q_{p_{ni}} = 0 = Q_{m_{ni}}$. The exchange matrices $\mathbf{Q}_m$ and $\mathbf{Q}_p$ become

$$\mathbf{Q}_p = \begin{pmatrix} Q_p & 0 \\ q_p^- & Q_p \end{pmatrix} \text{ and } \mathbf{Q}_m = \begin{pmatrix} Q_m & q_m^+ \\ 0 & Q_m \end{pmatrix},$$

Under strong competition (Eq. (12)), we deduce that the matrices $I_{nn}$ have negative coefficients. The **exclusion equilibrium with native hosts and symbiont is therefore stable**.

However, the matrix $I_{ii}$ has a negative coefficient

$$(1 - c_p/\mu_p)\, S^2 - Q_p Q_m/(\mu_p \mu_m)\,(1 - Q_{p_{ni}}/Q_p) < 0$$

and possibly a positive coefficient, if the symbiont is parasitic in the sense that

$$Q_{m_{in}} > \frac{c_m}{\mu_m} Q_m \;\Leftrightarrow\; \alpha_{ni} < \frac{q_{cm}}{q_{hm}} \beta_{in}\, d - \frac{c_m}{\mu_m} Q_m.$$

This inequality may be fulfilled if the symbiont is parasitic in the sense of definition of section B.2 and the invasive host is mutualistic enough to the symbiont. Otherwise, the native symbiont has to be really parasitic. As a result, when the native symbiont is parasitic enough relative to the invasive host, the equilibrium with **invasive species is unstable**.

**⑤ Invasive hosts are parasitic to native symbionts.** In this scenario, invasive hosts are parasitic to native symbionts, that is $Q_{p_{in}} = q_p^+ > Q_p > 0$ and $Q_{m_{in}} = q_m^- < 0$, while there is no association between native hosts and invasive symbionts, that is $Q_{p_{ni}} = 0 = Q_{m_{ni}}$. The exchange matrices $\mathbf{Q}_m$ and $\mathbf{Q}_p$ become

$$\mathbf{Q}_p = \begin{pmatrix} Q_p & 0 \\ q_p^+ & Q_p \end{pmatrix} \text{ and } \mathbf{Q}_m = \begin{pmatrix} Q_m & q_m^- \\ 0 & Q_m \end{pmatrix},$$

Under strong competition (Eq. (12)), the matrix $I_{ii}$ have negative coefficients. The **exclusion equilibrium with invasive hosts and symbiont is stable**.

However, the matrix $I_{nn}$ have a negative coefficient $Q_{m_{in}}/Q_m - c_m/\mu_m < 0$ and possibly a positive coefficient, if the host is really parasitic in the sense that

$$Q_{p_{in}} > Q_p \left( 1 + \left( \frac{c_p}{\mu_p} - 1 \right) \frac{S^2}{\frac{Q_p Q_m}{\mu_p \mu_m}} \right) \Leftrightarrow \beta_{in} < \frac{q_{hp}}{q_{cp}} \frac{\alpha_{ni}}{d} - Q_p \left( 1 + \left( \frac{c_p}{\mu_p} - 1 \right) \frac{S^2}{\frac{Q_p Q_m}{\mu_p \mu_m}} \right)$$

As a result, when the invasive host is parasitic enough relative to the native symbiont, the equilibrium with **native species is unstable**.

⑥ **Invasive symbionts are parasitic to native hosts.** This scenario is similar to the scenario ④. The **exclusion equilibrium with invasive species is stable** and if the invasive symbiont is parasitic enough relative to the native host, the equilibrium with **native species is unstable**.

⑦ **Native hosts are parasitic to invasive symbionts.** This scenario is similar to the scenario ⑤. The **exclusion equilibrium with native species is stable** and if the native host is parasitic enough relative to the invasive symbiont, the equilibrium with **invasive species is unstable**.

#### E.3 A simplified model on the proportion of native host and symbiont

In order to understand the outcome of the 7 scenario described in our manuscript, we will reformulate the problem by focusing on the proportion $\phi_p$ of native host and the proportion $\phi_m$ of native symbiont in the system, that are defined by

$$\phi_p = \frac{p_n}{p_n + p_i} \quad \text{and} \quad \phi_m = \frac{m_n}{m_n + m_i}$$

We also introduce the total biomass of the host $P = p_n + p_i$ and the total biomass of symbiont $M = m_n + m_i$. Let us remark that those quantities are related to $S$, $S = P/d + M$. Using the model (10), the proportions $\phi_p$ and $\phi_m$ satisfies the following dynamical system

$$\begin{cases} \phi_p' = \phi_p(1-\phi_p)\Big(\Big(\big(2Q_p - (Q_{p_{in}} + Q_{p_{ni}})\big)\phi_m - (Q_p - Q_{p_{ni}})\Big)\frac{M}{S} \\ \qquad\qquad\qquad\qquad\qquad\qquad + (c_p - \mu_p)(2\phi_p - 1)P\Big) \\ \phi_m' = \phi_m(1-\phi_m)\Big(\Big(\big(2Q_m - (Q_{m_{in}} + Q_{m_{ni}})\big)\phi_p - (Q_m - Q_{m_{in}})\Big)\frac{P}{S} \\ \qquad\qquad\qquad\qquad\qquad\qquad + (c_m - \mu_m)(2\phi_m - 1)M\Big) \end{cases} \tag{13}$$

Since $M$, $P$ and $S$ are not constant over time, the system is not autonomous and a simple phase plane won't explain the dynamics of $(\phi_p, \phi_m)$. However, we know from the full system (10), that those quantities are always positive and bounded. Thus, we can expect that the phase plane when we fix $M$, $P$ to a constant will explain the dynamics of the full system.

##### E.3.1 Steady states of the simplified system

Under the 7 scenarios (1)-(7), we assume that the competition between symbionts and hosts are strong ($c_m > \mu_m$ and $c_p > \mu_p$). Thus, the simplified model mainly contains trivial steady states corresponding to exclusion steady states of the full model

| | |
|---|---|
| $(1, 1)$ | native pair |
| $(0, 0)$ | invasive pair |
| $(1, 0)$ | mixed pair, invasive symbiont spillover |
| $(0, 1)$ | mixed pair, invasive host invasion |

Because the competition is strong, non trivial equilibrium will be unstable if they exist.

##### E.3.2 Phase plane of the simplified model for the 7 scenarios

The phase plane of the simplified model of Eq. (13) are provided in Fig. E.1, and can help us understand expected dynamics occurring in scenarios ①-⑦. A schematic representation of the outcomes of these scenarios is also provided in Fig. 3, in the main manuscript.

**① Native and invasive species do not share symbionts nor hosts.** In this scenario, we expect the equilibrium to reach either the ($p_n^*$, $m_n^*$,0,0) or the (0,0,$p_i^*$,$m_i^*$) steady state, as shown in Fig. E.1①. Which steady state is reached, depends on the initial conditions and model parameters, determining the size of the basin of attraction of each steady state. This situation of bistability is also discussed in scenario ① of the result section and illustrated in Fig. 3(a).

**② The association between native symbionts and invasive hosts is mutualistic.** Three steady states are possible, following the formation of novel mutualistic associations between invasive hosts and invasive symbionts, namely: ($p_n^*$,$m_n^*$,0,0), (0,$m_n^*$,$p_i^*$,0), and (0,0,$p_i^*$,$m_i^*$). A schematic representation of this scenario is provided in Fig. 3(b). Note that the steady state (0,$m_n^*$,$p_i^*$,0) can be reached through different routes, in orange and yellow in Fig. E.1②. A route in which the density of native symbionts is increased at first (in orange) and a route in which the density of invasive hosts is increased at first (in yellow). These two routes corresponds to the center left and center right pathways in Fig. 3(b).

**③ The association between native hosts and invasive symbionts is mutualistic.** Three steady states can be observed following the formation of novel associations between native hosts and invasive symbionts: ($p_n^*$,$m_n^*$,0,0), ($p_n^*$,0,0,$m_i^*$), and (0,0,$p_i^*$,$m_i^*$), where the steady state (0,$m_n^*$,$p_i^*$,0) can be reached through different routes, in orange and yellow in Fig. E.1③. A schematic representation of this scenario is provided in Fig. 3(c). The center left and right pathway in this figure correspond on the orange and yellow pathways in the phase plane. A route in which the density of native symbionts is increased at first (in orange) and a route in which the density of invasive hosts is increased at first (in yellow). These two routes corresponds to the center left and center right pathways in Fig. 3(b).

**④ Native symbionts are parasitic to invasive hosts, and ⑦ Native hosts are parasitic to invasive symbionts.** In these cases, only one steady state ($p_n^*$,$m_n^*$, $0, 0$) is stable, as the exclusion of invasive species (either hosts or symbionts) leads to the extinction of their invasive partner, as seen in Fig. E.1④ and ⑦ and in Fig. 3(a).

**⑤ Invasive hosts are parasitic to native symbionts, and ⑥ Invasive symbionts are parasitic to native hosts.** In these cases, the only stable steady state is (0,0,$p_i^*$,$m_i^*$), as shown in Fig. E.1⑤ and ⑥ and in Fig. 3(a). This is because the exclusion of native hosts or symbionts, due to exploitation by invasive species, leads as well to the exclusion of the native mutualistic partner.

### F Multiple hosts and multiple symbionts model

Our framework can be extended to consider multiple hosts and symbionts. If we consider $M$ competing hosts exchanging resources with $N$ competing symbionts, we obtain:

$$\begin{cases} \dfrac{dp_j}{dt} &= r_{p_j} p_j + \dfrac{p_j}{\dfrac{\sum_j p_j}{d} + \sum_i m_i} \sum_i m_i Q_{p_{ij}} - \mu_p p_j^2 - c_p \sum_{k \neq j} p_k \,, \quad \text{for} \quad j = 1, ..., M \,, \\ \dfrac{dm_i}{dt} &= \dfrac{m_i}{\dfrac{\sum_j p_j}{d} + \sum_i m_i} \sum_j p_j Q_{m_{ji}} - \mu_m m_i^2 - c_m \sum_{l \neq i} m_l \,, \quad \text{for} \quad i = 1, ..., N \,, \end{cases}$$

where $Q_{p_{ij}}$ is the effect of symbiont $i$ on host $j$ and $Q_{m_{ji}}$ is the effect of host $j$ on symbiont $i$. These are defined for any $i \in \{1, \ldots, N\}$ and $j \in \{1, \ldots, M\}$ by

$$Q_{p_{ij}} = q_{hp} \frac{\alpha_{ji}}{d} - q_{cp} \beta_{ij} \quad \text{and} \quad Q_{m_{ij}} = q_{cm} \beta_{ji} - q_{hm} \frac{\alpha_{ij}}{d} \,.$$

An analysis of this system is beyond the scope of this paper, and will be left to future work.

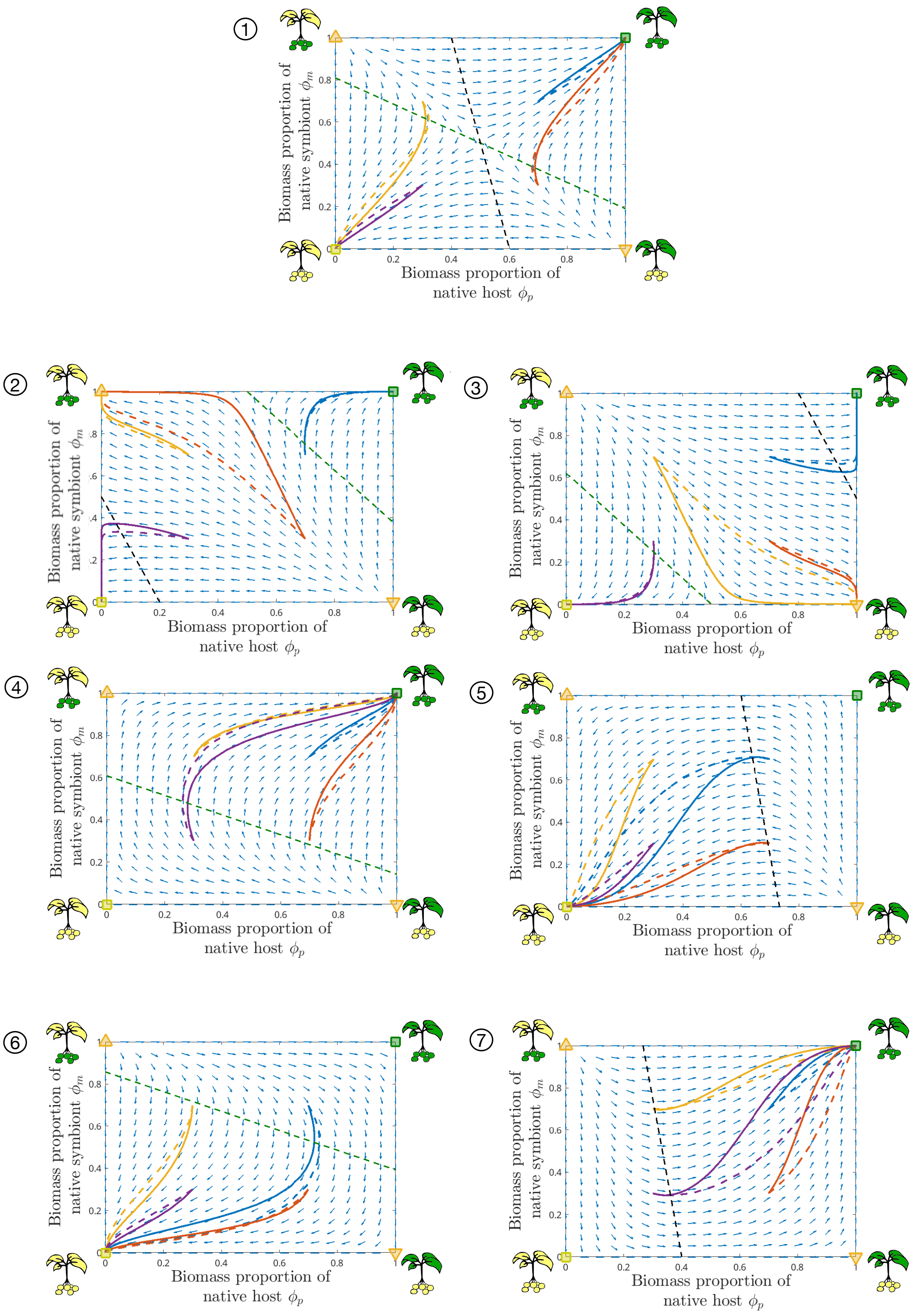

**Fig. E.1:** Phase plane of the approximation model (13). The green square corresponds to the exclusion equilibrium with native species, the yellow square corresponds to the exclusion equilibrium with invasive species, and the orange triangles correspond to mixed pair equilibria. Shown are 4 trajectories of the full system (10) (solid curves) and the approximated model (dashed curves). The straight dashed lines correspond to the nullclines of the system, and the arrows are the approximation of the flow (13).
